# Single-cell chromatin tracing reveals multimodal molecular programs during memory formation

**DOI:** 10.64898/2026.06.16.732522

**Authors:** Kaho Itoh, Valentina Khalil, Islam Faress, Taro Kitazawa

## Abstract

Experience-dependent activity is converted to coordinated molecular programs in neuronal ensembles during memory formation. However, due to the sparsity of the ensembles and the transience of immediate early gene (IEG) expression, it is unclear how IEGs engage downstream secondary response genes (SRGs) to regulate learning-specific neuroplasticity. Here, we generated a single-cell multiomic atlas of aversive learning and developed ChromTRAP, which retrospectively identifies recently activated neuronal ensembles from AP-1 (FOS/JUN)-centred chromatin traces. We integrated transcription, chromatin accessibility, histone modifications, and FOS occupancy across the amygdala, hippocampus, and prefrontal cortex. This revealed regulatory programs of learning-associated genes (LAGs), defined as SRGs preferentially induced by associative learning relative to baseline activity or independent stimulus exposure. These programs followed a brain-region- and cell-type-specific proximal-distal regulatory logic: gene-proximal Polycomb-associated H3K27me3 remodeling and AP-1-bound H3K27ac-marked distal enhancers. LAGs were further associated with enhanced intercellular signaling and MEF-family activity. Our findings establish a single-cell multiomic framework for linking learning experience to layered epigenetic regulation during activity-dependent neuroplasticity.

## Introduction

Memory is allocated to sparse neuronal ensembles, termed memory engrams that are activated during learning and reactivated during recall across multiple brain regions ^1–5^. This process triggers the rapid induction of immediate early genes (IEGs; e.g., *Fos*), many of which encode transcription factors. These IEGs drive downstream secondary response genes (SRGs) that serve as the effectors of plasticity ^6^, through regulation of multilayered epigenomic landscapes (e.g., chromatin accessibility, histone modifications, DNA methylation, chromatin loops) ^7–11^ . However, neuronal activation and IEG expression also occur under baseline conditions. It remains unclear how IEGs selectively engage learning-associated SRG programs, rather than nonspecific activity-induced transcription, during memory formation.

A major barrier to addressing this issue has been the difficulty of identifying the relevant neuronal ensembles during the temporal window that links IEG induction to SRG activation. To date, putative engram cells have been operationally defined and isolated using molecular marker-based approaches, including antibody-based fluorescence-activated cell sorting (FACS) against IEG proteins ^12,13^ or genetically encoded activity reporters such as Targeted Recombination in Active Populations (TRAP; e.g., *Fos::CreERT2*) ^5,10,14–19^. These approaches capture different temporal windows after learning: FOS-antibody-based strategies enrich for neurons within ∼1 h of activation, whereas genetic recorders require substantial time, ranging from ∼12 h to days, to label activated ensembles. Critically, during the intermediate window when SRG expression becomes prominent, approximately 4-6 h after learning, IEG mRNA and protein have largely declined, while reporter-based strategies have not yet reached detectable levels. As a result, the genome-wide SRG programs and regulatory mechanisms operating in sparse neuronal ensembles remain poorly understood.

A further challenge is the heterogeneity of learning-associated transcriptional and epigenomic programs, as they vary across brain regions and neuronal types ^20,21^. Neuronal ensembles are distributed across brain regions, including the amygdala (e.g., lateral amygdala, LA; basal amygdala, BA), hippocampus (e.g., CA1, CA3, DG), and medial prefrontal cortex (mPFC), each of which contributes distinctly to memory processing ^1,21^.

Moreover, recent studies indicate that engram formation and function are shaped by neuronal type, highlighting the need to resolve ensemble programs at both regional and cell-type levels ^20,22^. However, in contrast to bulk-level analyses, single-cell epigenomic profiling of these ensembles remains at an early stage, leaving how cell-type- and region-specific chromatin states coordinate gene regulatory programs in neuronal ensembles unresolved.

To overcome these limitations, we generated a single-cell multiomic atlas of aversive learning and developed Chromatin-based Targeted Recovery of Activated Populations (ChromTRAP) (Figure 1A). ChromTRAP is a computational approach for retrospectively identifying cells with recent IEG-expression history during the SRG expression window from single-cell multiomics chromatin traces. We integrated ChromTRAP with a combination of single-cell multiomic approaches, including established epigenetic platforms and a newly developed method for transcription factor binding analysis (i.e., multimodal nanobody-mediated CUT&Tag for transcription factor; multi-nanoCTF). Leveraging these approaches, we jointly profiled transcription, chromatin accessibility, and histone modifications during aversive memory formation across basolateral amygdala (BLA, BA + LA), dorsal hippocampus (dHPC), and mPFC. Furthermore, we used chemogenetically induced FOS multi-nanoCTF to validate AP-1 occupancy at activity-regulated enhancers. Our study provides an integrated framework linking activity-associated chromatin traces with chromatin state, enhancer selection, intercellular signalling, and learning-specific transcription. We revealed how memory-associated neuronal ensembles engage coordinated transcriptional and epigenomic programs during memory formation.

**Figure 1.**
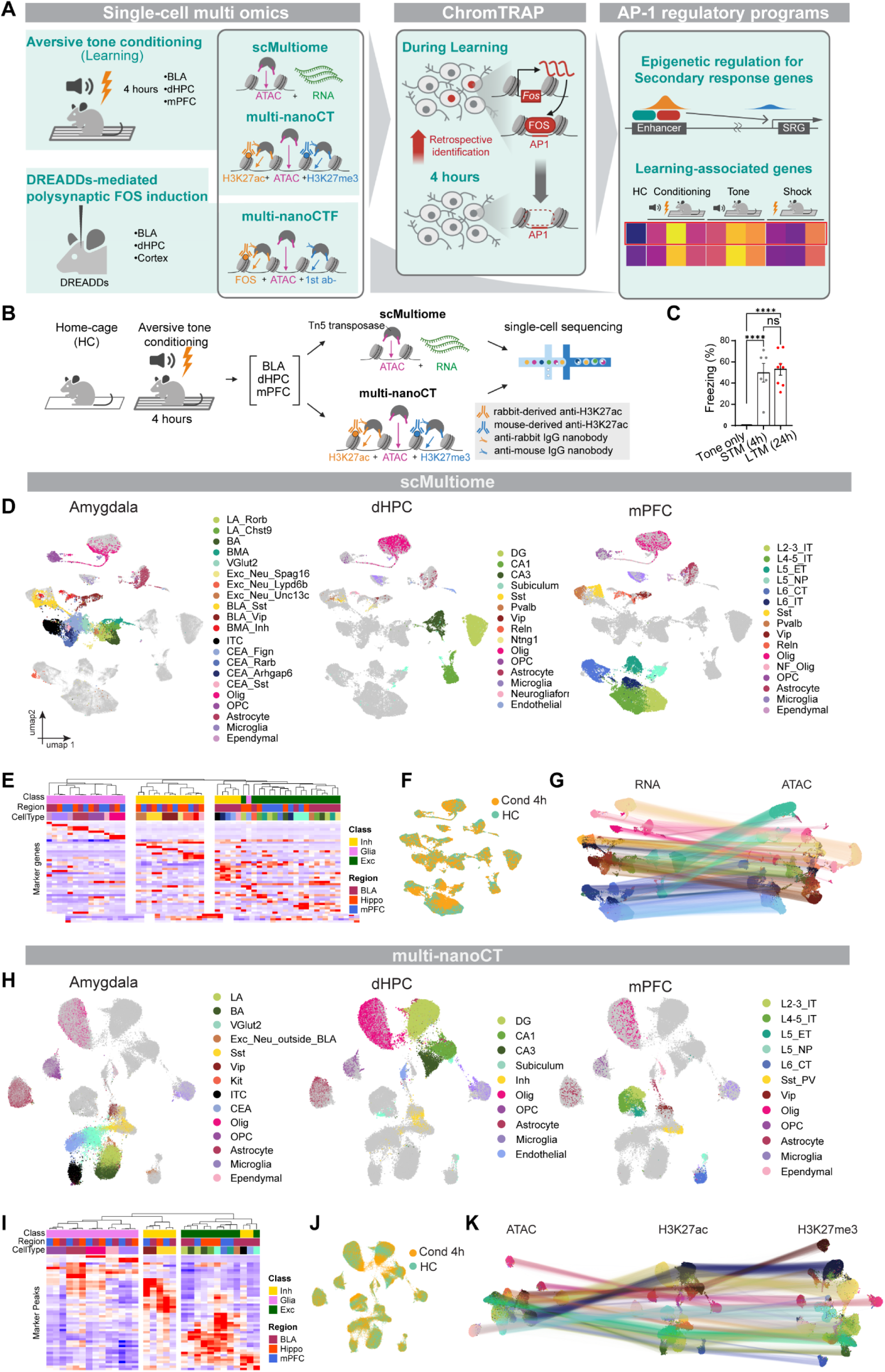
Multiomics single-cell profiling during memory formation. (A) Overview of this study. Left: Transcriptome, chromatin accessibility, and histone modifications were profiled at single-cell resolution 4 h after aversive tone conditioning using single-cell Multiome (scMultiome) and multi-nanobody-mediated CUT&Tag (multi-nanoCT). Single-cell FOS occupancy was validated after DREADDs (Designer Receptors Exclusively Activated by Designer Drugs)-mediated brain-wide activation using multi-nanobody-mediated CUT&Tag for transcription factors (multi-nanoCTF). Middle: Neuronal ensembles with recent IEG-expression history were retrospectively identified using the ChromTRAP computational framework. Right: AP-1 regulatory programs of learning-associated genes (LAGs) were analysed. (B) Experiment scheme for scMultiome (83,611 cells) and multi-nanoCT (123,876 cells) from home-cage (HC) and conditioned animals. scMultiome jointly profiles transcriptome and chromatin accessibility (ATAC). multi-nanoCT combined ATAC tagmentation with modality-barcoded nanobody–Tn5 targeting species-specific primary antibodies, enabling ATAC, H3K27ac, and H3K27me3 profiling from the same nuclei. (C) Freezing during recall in tone-only and conditioned mice. STM, short-term memory recall 4 h after conditioning; LTM, long-term memory recall 24 h after conditioning. Statistical significance was assessed by one-way ANOVA followed by Tukey’s multiple-comparisons test. ns, not significant; *P < 0.05; **P < 0.01; ***P < 0.001; ****P < 0.0001. (D, H) UMAPs derived from scMultiome (D) and multi-nanoCT (H). Cell-type representations based on RNA (scMultiome) (D) and H3K27ac (multi-nanoCT) (H) modalities, respectively. (E, I) Heatmaps show gene expression profiles of selected marker genes in the scMultiome dataset (E) and H3K27ac level of cell-type marker peaks in the multi-nanoCT dataset (I). Rows represent marker genes or peaks, and columns represent annotated cell types from BLA, hippocampus, and mPFC. Heatmap colours indicate row-wise z-scored mean expression or accessibility. Columns were hierarchically clustered using average linkage based on Pearson correlation distance of row-wise z-scored marker profiles. (F, J) Representation of experimental conditions. (G, K) Representation of modalities. Lines connect matched modalities measured from the same nuclei, illustrating cross-modality correspondence within individual cells. Abbreviations for cell types: BA, basal amygdala; CEA, central amygdala; CT, corticothalamic-projecting; ET, extratelencephalic-projecting; Exc, excitatory neurons; Inh, inhibitory neurons; IT, intratelencephalic-projecting; ITC, intercalated cells; L, cortical layer; LA, lateral amygdala; NF_Olig, newly formed oligodendrocyte; NP, near-projecting; Olig, oligodendrocyte; OPC, oligodendrocyte progenitor cells.

## Results

### Multimodal single-cell profiling of memory formation across brain regions

To characterize molecular states during memory formation, we subjected mice to aversive tone conditioning and microdissected the BLA, dHPC, and mPFC 4 h after conditioning for single-cell multiomics analysis (Figures 1A and 1B). To validate the efficacy of the behavioural paradigm, independent cohorts were evaluated for recall at 4 h and 24 h post-conditioning. At both time points, conditioned mice exhibited tone-evoked increased freezing relative to tone-only controls, confirming successful memory acquisition and retrieval (Figures 1C and S1A).

We first performed 10x Genomics Single Cell Multiome ATAC + Gene Expression assay (scMultiome) to jointly profile gene expression and chromatin accessibility (i.e., ATAC profile), in the same nuclei (Figure 1B). To extend this analysis to histone modifications, we next performed nanobody-mediated single-cell CUT&Tag (nanoCT) ^23,24^ to simultaneously profile three modalities in the same nuclei: chromatin accessibility, transcriptionally active H3K27ac mark, and Polycomb-repressive H3K27me3 histone mark (Figure 1B). Rather than applying nanoCT separately to each sample, we adapted the sample-pooling strategy we previously established for scHisTrac-seq to nanoCT, generating a multiplexed workflow termed multi-nanoCT. In this approach, multiple samples are loaded into a single 10x microfluidic reaction to reduce cost and minimize batch effects while preserving modality-specific chromatin readouts (Methods) ^25^.

To ensure high-fidelity single-cell data, stringent quality control criteria were applied to remove doublets, dead cells, and low-quality nuclei (Methods). Following this quality control, we retained 83,611 cells from scMultiome (BLA 23,151; dHPC 24,694; mPFC 35,766) and 123,876 cells from multi-nanoCT (BLA 47,385; dHPC 50,699; mPFC 25,792), derived from two biological replicates per experimental condition and brain region (Figures 1D-1K and S1B). The quality-control metrics were consistent with those reported in the original studies (Figures S1C and S1D) ^23,26^. We clustered scMultiome and multi-nanoCT cells based on mRNA and H3K27ac, respectively (Figures 1D and 1H). scMultiome clusters were annotated using canonical cell-type-marker genes (Figure 1E; Table S1). To annotate multi-nanoCT clusters, we used the annotated scMultiome dataset as a reference: cell-type-specific ATAC peak sets were derived from scMultiome and then used to assign multi-nanoCT cell types based on H3K27ac enrichment (Figure 1I; Table S2). In both datasets, cells segregated primarily by cell type, whereas samples from home-cage and conditioned animals were distributed uniformly across the same cell-type clusters (Figures 1D, 1F, 1H, and 1J), indicating that cell type identity is the dominant source of variation relative to learning-induced activity state.

Across all brain regions, spatially and molecularly distinct populations of excitatory neurons were identified: two subtypes of LA and one of BA in the amygdala, hippocampal DG, CA1 and CA3, and specific cortical layers of mPFC. These populations clustered independently, suggesting different regional molecular identities. In contrast, inhibitory neurons (e.g., Sst, Vip) and glial cells (e.g., astrocyte and oligodendrocyte) from different brain regions clustered together, indicating highly conserved molecular features across brain areas (Figures 1D, 1E, 1H, 1I).

We next clustered cells using the remaining individual modalities, namely, ATAC for scMultiome and H3K27me3 and ATAC for multi-nanoCT (Figures 1G and 1K). These modalities recapitulated the same major cell-type structure on UMAP, indicating that cell-type identity can be robustly resolved across assay modalities.

Together, these datasets provide a multimodal single-cell resource for profiling transcriptional and epigenomic states across memory-related brain regions and cell types during memory formation.

### ChromTRAP retrospectively identifies activated neurons through chromatin traces

Next, to characterize gene regulatory programs in activated neuronal ensembles, we developed ChromTRAP, a computational framework to identify activated neurons from single-cell multiomics data. At 4 h after conditioning, when SRG expression becomes prominent, *Fos* mRNA and FOS protein have largely declined, limiting their utility as activity markers (Figures 2A left, 2B, S2A, and S2B top). We therefore established an alternative strategy to retrospectively identify neurons with recent IEG expression history using persistent chromatin traces (Figure 2C).

**Figure 2.**
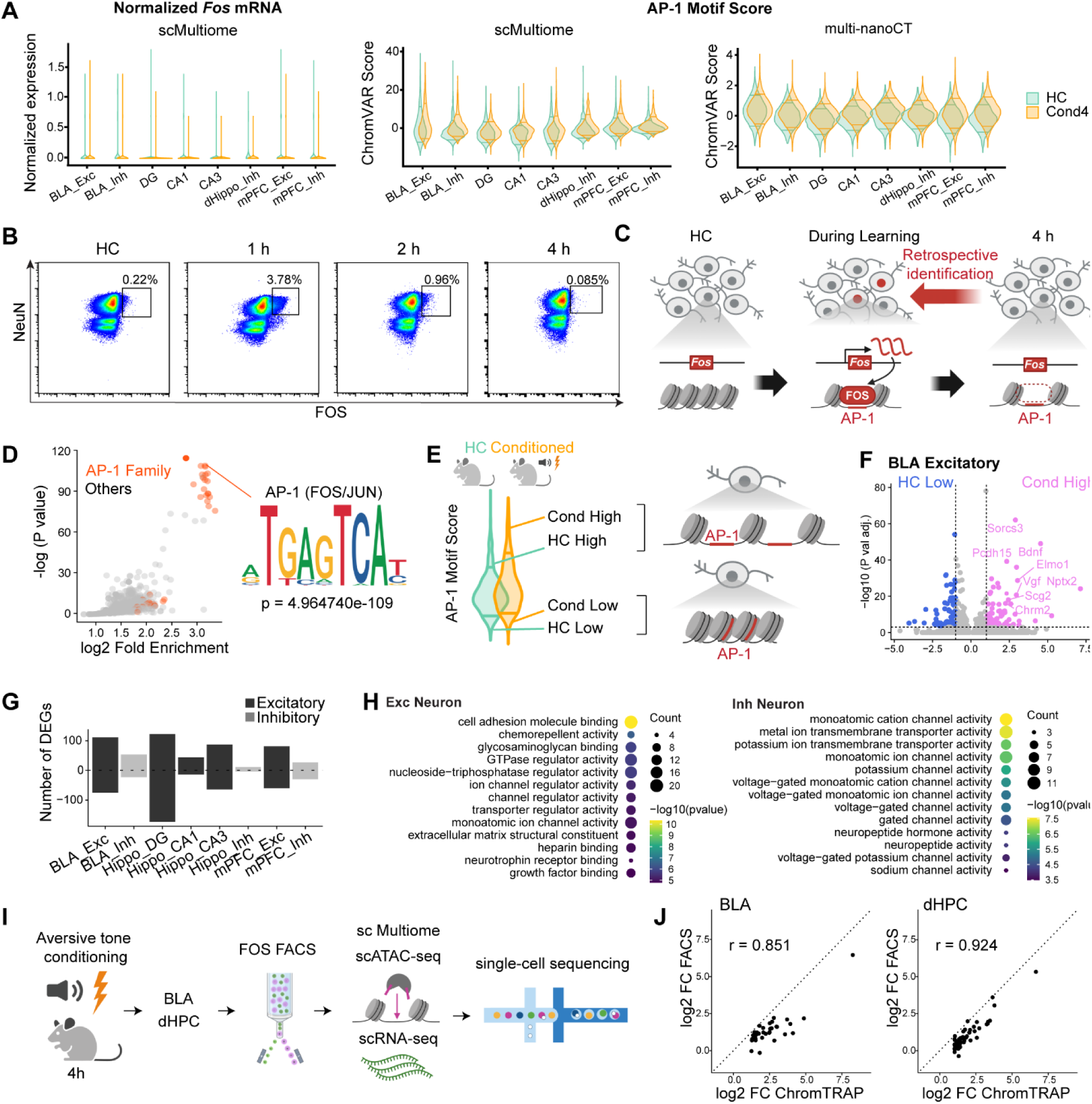
ChromTRAP retrospectively identifies activated neurons using chromatin traces. (A) *Left:* Log2-normalized *Fos* mRNA levels in home-cage (HC) mice and 4 h after conditioning (Cond4h) across cell types and brain regions. *Right:* AP-1 Motif scores (i.e., chromVAR deviation scores) quantified based on ATAC signals in scMultiome and multi-nanoCT. The horizontal lines indicate the 10th and 90th percentiles. The 90^th^ percentiles were shifted upward in Cond4h neurons compared with HC neurons. (B) Representative FOS-FACS profiles from BLA in HC mice and at 1 h, 2 h, or 4 h after conditioning. NeuN was also used to enrich neurons. (C) Scheme of ChromTRAP. AP-1 (FOS/JUN)-centred chromatin traces are used to retrospectively infer recent IEG-expression history after *Fos* mRNA and protein levels have declined. (D) Volcano plot showing transcription factor motif enrichment in Cond-gained peaks from BLA excitatory neurons relative to matched background peaks. AP-1-related motifs are shown in orange, and other motifs are shown in grey. (E) Schematic of ChromTRAP-based neuron stratification. Top-scoring conditioned neurons ranked by AP-1 motif score were defined as Cond High, whereas top-scoring HC neurons were defined as HC High and low-scoring neurons as Cond Low or HC Low. (F) Representative volcano plots from DEG comparisons, showing Cond High versus HC Low neurons in BLA excitatory population. (G) Number of significantly differentially expressed genes identified across cell types and brain regions. Positive and negative values indicate up- and down-regulated genes, respectively. (H) GO terms enriched in upregulated genes after aggregation across brain regions for excitatory and inhibitory neurons. (I) Experimental scheme for benchmarking ChromTRAP using FOS protein-FACS isolation followed by scMultiome profiling. (J) Scatter plot comparing log2 fold changes of ChromTRAP-derived DEGs between ChromTRAP-defined Cond High neurons and FOS-FACS-isolated neurons. Pearson’s correlation is shown. Concordant directions and larger fold changes in ChromTRAP-defined neurons support improved sensitivity for detecting activity-associated DEGs during the SRG window.

Because AP-1 complexes (e.g. FOS, FOSB, and JUNB) are induced by neuronal activity and bind activity-regulated enhancers that remain accessible for hours after AP-1 expression declines ^6–8^, we hypothesized that chromatin accessibility at AP-1 sites could serve as a molecular trace of recent activation. Supporting this, unbiased motif analysis revealed broad enrichment of AP-1 family motifs in chromatin regions that gained accessibility 4 h after conditioning (Figure 2D).

Based on this principle, we developed ChromTRAP, a framework that utilizes chromVAR ^27^ to quantify a per-cell deviation score for the MA1141.2 AP-1 motif (“AP-1 motif score”) in the ATAC modality of single-cell multiomics data. While *Fos* mRNA levels failed to separate between home-cage and conditioned samples at 4 h after conditioning (Figures 2A and S2B top), the AP-1 motif score clearly distinguished these samples across brain regions and cell types (Figures 2A right and S2B middle and bottom). This indicates that AP-1-associated accessibility provides a more robust basis for identifying recent IEG expression during the SRG expression window.

We therefore used AP-1 motif scores to stratify neurons into activity-associated populations: top- and low-scoring neurons in conditioned samples were designated “Cond High” and “Cond Low,” respectively, while the corresponding populations in home-cage (HC) samples were designated “HC High” and “HC Low” (Fig. 2E; Methods). This classification enabled comparison of activated neuronal ensembles with baseline ensembles without relying on contemporaneous *Fos* expression.

### Benchmarking of ChromTRAP

We next carried out benchmarking of ChromTRAP. First, to assess whether ChromTRAP-defined neuronal ensembles reflect neuronal activation beyond AP-1 activity alone, we examined correlations between chromVAR scores for the AP-1 motif and other transcription factor motifs (Figure S2C). We observed strong positive correlations with neuronal activity-related motifs including CREB-, PAS-domain factors (e.g., NPAS family), NR4A, and EGR. This suggests that ChromTRAP-defined neuronal ensembles engage broader activity-dependent regulatory programs. This is consistent with previous reports showing increased overlap between FOS- and NPAS4-expressing neuronal populations following aversive conditioning ^28^.

To analyze activity-response programs in ChromTRAP-defined neuronal ensembles, we identified differentially expressed genes (DEGs) as genes significantly upregulated in any of three comparisons: Cond High versus Cond Low, Cond High versus HC Low, and HC High versus HC Low (Table S3). This analysis identified a larger number of DEGs in excitatory than inhibitory neurons (Figures 2F and 2G). Within the dHPC, the DG showed the largest number of DEGs, followed by other pyramidal neurons, whereas inhibitory neurons showed the fewest, consistent with a previous report ^12^. Across brain regions, up-regulated genes included established markers of neuronal plasticity, such as *Bdnf, Sorcs3, Nptx2* and *Scg2* ^29–32^. Gene ontology analysis further revealed enrichment of cell–cell interaction-related functions in DEGs from both excitatory and inhibitory neurons, including cell adhesion molecule binding and potassium ion transmembrane transporter activity (Figures 2H and S2D).

We further tested whether ChromTRAP-defined DEG detection was robust to baseline selection by stratifying neurons into finer AP-1 motif score bins. ChromTRAP-associated DEGs increased progressively with AP-1 motif score and peaked in the top-scoring population (Figure S2E), indicating that top-scoring neurons retain a clear transcriptional contrast even when intermediate-score neurons, rather than HC Low or Cond Low neurons, are used as the baseline. Thus, ChromTRAP-defined DEG detection is robust to the choice of baseline population.

We next evaluated ChromTRAP-defined DEGs using a FOS protein-based FACS strategy, which provides direct protein-level enrichment of recently activated neurons. We applied this approach 4 h after conditioning to match the SRG time window interrogated by ChromTRAP. Because FOS protein has substantially declined at this time point (Figure 2B),

FOS-FACS is expected to recover only a subset of neurons retaining detectable FOS signal. This provides a complementary, conservative protein-based benchmark for activity-associated DEG detection during SRG window.

We recovered FOS-positive neurons from the BLA and dHPC by FACS and performed scMultiome profiling (Figure 2I). This low FOS-positive fraction constrained the number of nuclei for 10x Chromium loading (Figures S3A-C and S3H). Consequently, the single-cell recovery rate after quality control was only 25% for FOS-FACS samples (1,631 and 1,672 cells in BLA and dHPC, respectively), compared with 63.1% for density-centrifuged samples processed without rare-population sorting, indicating that rare-population isolation reduces recovery efficiency and data yield after single-cell sequencing (Table S4). After cell-type identification (Figures S3D and S3I), we compared DEGs identified from ChromTRAP-defined Cond High neurons and FOS-FACS-isolated neurons against matched controls.

ChromTRAP identified substantially more DEGs than FOS-FACS in both the BLA (35 versus 15) and dHPC (58 versus 8) (Figures S3E-G and S3J-L). Despite this difference in sensitivity, the two approaches showed concordant transcriptional changes: a subset of DEGs overlapped, and log2 fold change of DEGs were positively correlated (Pearson’s r = 0.851 in the BLA and 0.924 in the dHPC) (Figure 2J). Moreover, most DEGs exhibited larger effect sizes in ChromTRAP dataset. These results indicate that ChromTRAP captures transcriptional signatures similar to those identified by FOS-FACS-based experiments, while providing improved sensitivity for detecting DEGs.

Together, these data establish ChromTRAP as an effective strategy for retrospectively identifying neurons with recent IEG-expression history during the SRG expression window. By defining neuronal ensembles from the ATAC modality, ChromTRAP enables integrated analysis of matched gene-expression, histone-modification, and chromatin-accessibility layers within the same single-cell multiomic datasets.

### Polycomb-chromatin remodeling in activity-dependent transcriptional programs

Having identified ChromTRAP-defined activity-responsive transcriptional programs, we asked whether these DEGs were linked to gene-proximal Polycomb-associated chromatin remodeling. Polycomb-repressive H3K27me3 mark has been reported to regulate activity-dependent expression of *Bdnf* ^33^. Furthermore, a previous study has comprehensively characterized the remodelling of H3K27me3 mark in activity-dependent transcription during development^34^. However, whether Polycomb-associated chromatin remodeling broadly contributes to learning-induced transcriptional programs in the adult brain remains unresolved. We therefore systematically examined Polycomb-H3K27me3 dynamics during memory formation by defining neuronal ensembles with ChromTRAP and analysing the matched H3K27me3 modality in multi-nanoCT datasets.

We first defined Polycomb-covered genes as those overlapping H3K27me3 domains. Across brain regions and neuronal types, more than half of ChromTRAP-based DEGs were Polycomb-covered (Figure 3A). We next examined conditioning-associated H3K27me3 and chromatin accessibility dynamics at these DEGs by comparing ChromTRAP-Cond High against Cond Low neurons. This revealed that DEGs were significantly enriched in genomic regions with reduced H3K27me3 and increased ATAC accessibility compared with non-DEG controls (Figure 3B). Direct signal comparison within DEG-overlapping H3K27me3 broad peaks confirmed this trend, revealing higher ATAC accessibility and lower H3K27me3 in Cond High neurons than in Cond Low neurons (Figure S4A and S4B).

**Figure 3.**
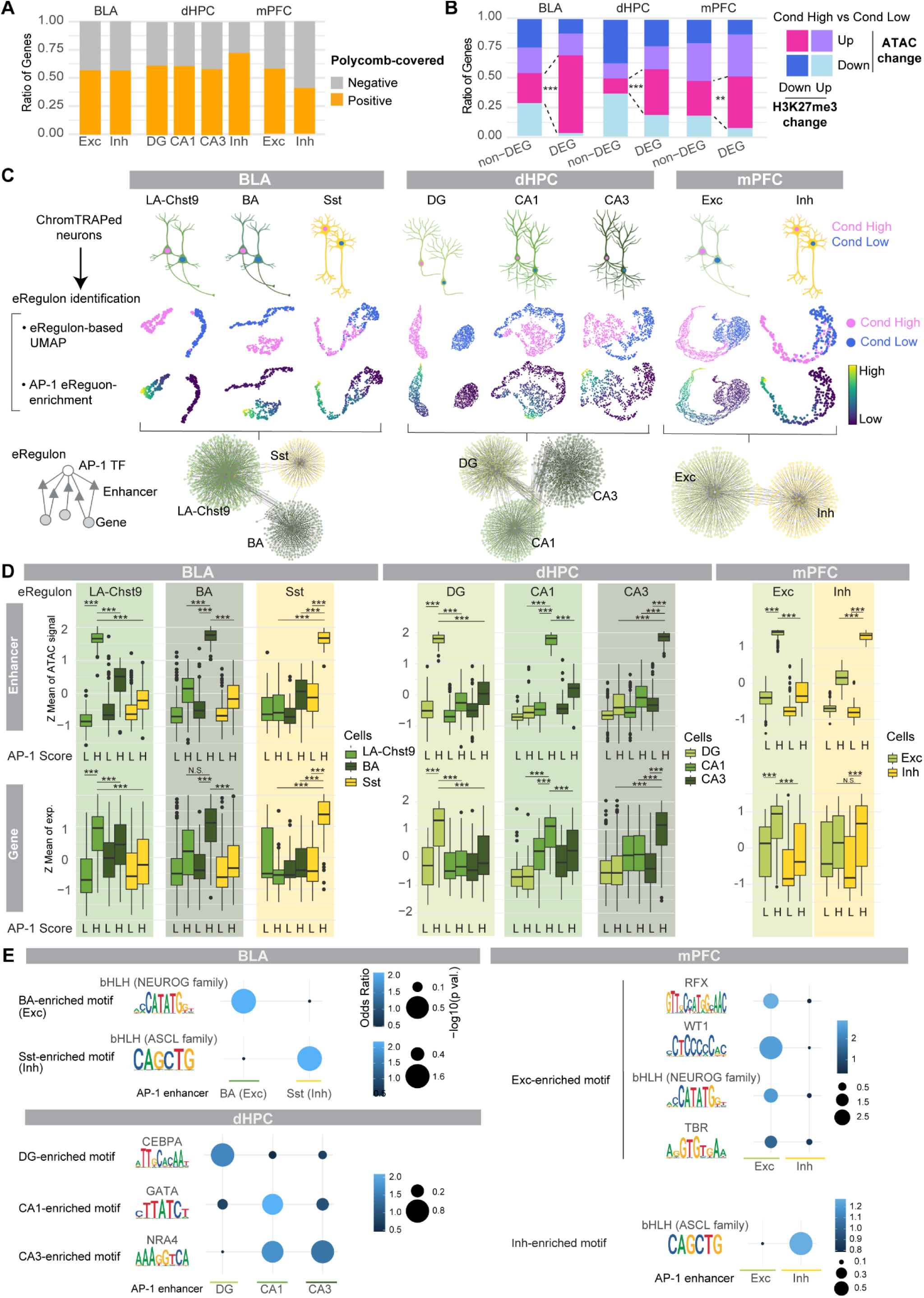
Epigenetic regulation of activity-response genes. (A) Bar plots showing the fraction of ChromTRAP-defined DEGs overlapping Polycomb-H3K27me3-covered genomic regions across neuronal cell types and brain regions. (B) Bar plots showing activity-associated epigenetic changes in Polycomb-H3K27me3 domains. DEGs and non-DEGs were classified by the direction of H3K27me3 and ATAC fold changes between Cond High and Cond Low neurons. DEGs were enriched for loci showing increased ATAC accessibility and reduced H3K27me3 in Cond High neurons. Excitatory neurons are shown. Statistical significance of differences in category distributions between DEGs and non-DEGs were assessed using the two-sided Fisher’s exact tests. ***P < 0.001. (C) Overview of ChromTRAP-SCENIC+-based gene regulatory network analysis. SCENIC+ was used to infer enhancer-driven regulons (eRegulons), defined here as transcription factor-associated enhancer–target gene modules. Top, ChromTRAP-defined Cond High neurons were compared with Cond Low neurons for each cell type across brain regions. Middle, eRegulon-based UMAP embeddings for each cell type, showing separation of Cond High and Cond Low neurons and AP-1 eRegulon enrichment across examined cell types. Bottom, Network plots showing AP-1 eRegulon-associated target genes and enhancers across neuronal cell types. Only cell types in which AP-1 eRegulons were robustly identified are shown (Methods). (D) Box plots showing ATAC accessibility (top) and target-gene expression (bottom) for cell-type-specific AP-1 eRegulons. Signals are compared between Cond High (AP-1 Score H) and Cond Low (AP-1 Score L) neurons within the matched cell type, showing increased enhancer accessibility and target-gene expression in Cond High neurons. Statistical significance was assessed using CAMERA implemented in limma. *P < 0.05; **P < 0.01; ***P < 0.001. (E) Bubble plots showing enrichment of cell-type-specific transcription factor motifs in AP-1 eRegulon-associated enhancers. Dot colour indicates odds ratio, and dot size indicates statistical significance. P values were calculated using one-sided Fisher’s exact tests.

Together, these results reveal a gene-proximal Polycomb-H3K27me3 regulatory layer associated with activity-responsive transcription in ChromTRAP-defined neuronal ensembles.

### Cell-type-specific AP-1 enhancer-gene regulatory networks

Following the analysis of gene-proximal Polycomb regulation, we next turned to distal enhancer regulation during memory formation. AP-1 is a major regulator of neuronal activity-dependent transcription and drives downstream SRG expression by binding activity-regulated distal enhancers ^8^. This was supported by our unbiased motif screening (Figure 2D). However, how AP-1-associated enhancer–gene networks are organized within sparse neuronal ensembles, and how these programs differ across brain regions and neuronal cell types, remains poorly understood. We therefore addressed this by integrating ChromTRAP with SCENIC+ ^35^, a framework that utilizes single-cell gene expression and chromatin accessibility profiles to infer enhancer-driven regulon (eRegulon), which is an integrated regulatory unit comprising a transcription factor and its coordinate target enhancers and genes (Figure 3C).

We applied the SCENIC+ workflow to Cond High and Cond Low populations of the scMultiome dataset. To reduce the dominant effect of cell-type differences on gene regulatory network inference, we performed the analysis separately for each neuronal type (Figure 3C, top, see Methods). mPFC excitatory neurons were analysed jointly across layers, as this did not introduce significant cell-type-driven biases. Region-based eRegulon UMAPs clearly separated Cond High vs Low neurons (Figure 3C, middle), confirming that the inferred eRegulons captured activity-associated regulatory states. We identified “AP-1 eRegulons” across all examined cell types (Figure 3C, middle).

The target genes and associated enhancers of AP-1 eRegulons showed limited overlaps across cell types, indicating that AP-1-associated regulatory networks are highly cell-type-specific (Figure 3C, bottom). To more rigorously define cell-type-specific AP-1 regulatory programs, we divided AP-1 eRegulons into shared and cell-type-specific groups (Figure S4C; Table S5, see Methods). We confirmed that cell-type-specific AP-1 eRegulons displayed elevated accessibility in their matched cell types (Figure 3D, top). Furthermore, these eRegulons were also accompanied by concordant target-gene expression patterns (Figure 3D, bottom).

We next asked how the cell-type specificity of these AP-1-associated enhancers is established. A preceding study has shown that AP-1-bound regulatory elements are co-specified by cell-type-specific transcription factors ^36^. We therefore tested whether a similar principle applies across neuronal types by screening neuronal type-specific transcription factors using chromVAR and quantifying occurrence of their binding motifs in AP-1 eRegulon-associated enhancers. We identified transcription factor motifs characteristic of each neuronal type that were significantly overrepresented in the corresponding AP-1 eRegulon-associated enhancers (Figures 3E and S4D). For instance, in BLA and mPFC, excitatory and inhibitory neuron-specific enhancers were enriched for the NEUROG and ASCL bHLH motifs, respectively, consistent with their roles as corresponding lineage-defining regulators ^37,38^. These findings support a model in which AP-1-regulatory landscapes are shaped by cell-type-specific transcription factors even among closely related neuronal types.

Lastly, because AP-1 eRegulon-associated enhancers showed limited overlap with Polycomb-repressive H3K27me3 (Figure S4E), we reasoned that activity-dependent distal enhancer regulation is likely mediated by regulatory mechanisms distinct from gene-proximal Polycomb remodelling.

Together, these analyses suggested a distal enhancer regulatory layer in which AP-1-associated enhancer–gene programs are specified by cell-type-specific transcription factor landscapes during memory formation.

### FOS occupancy links AP-1 to active enhancer states

Because the AP-1 eRegulons described above were inferred from chromatin accessibility and linked gene expression, two limitations remained. First, these analyses did not directly establish whether AP-1/FOS physically occupies the predicted enhancers. Second, increased accessibility can indicate regulatory potential or enhancer priming, but does not by itself demonstrate acquisition of an active enhancer state: H3K27ac serves as a direct marker of transcriptionally active enhancers ^39^. Thus, we sought to directly measure FOS occupancy at predicted AP-1 enhancers and determine whether this binding is coupled to H3K27ac-marked enhancer activation.

To address this, we developed multiplexed nanobody-mediated single-cell CUT&Tag for transcription factor (multi-nanoCTF), a method for single-cell profiling of transcription factor occupancy. Multi-nanoCTF extends the multi-nanoCT strategy by enabling transcription factor binding, chromatin accessibility, and matched 1st antibody-minus control in the same nuclei (Figures 4A, S5A-C). The 1st antibody-minus modality served as an essential internal control for non-specific Tn5-derived background, which often correlates with chromatin accessibility and can confound detection of true transcription factor occupancy.

**Figure 4.**
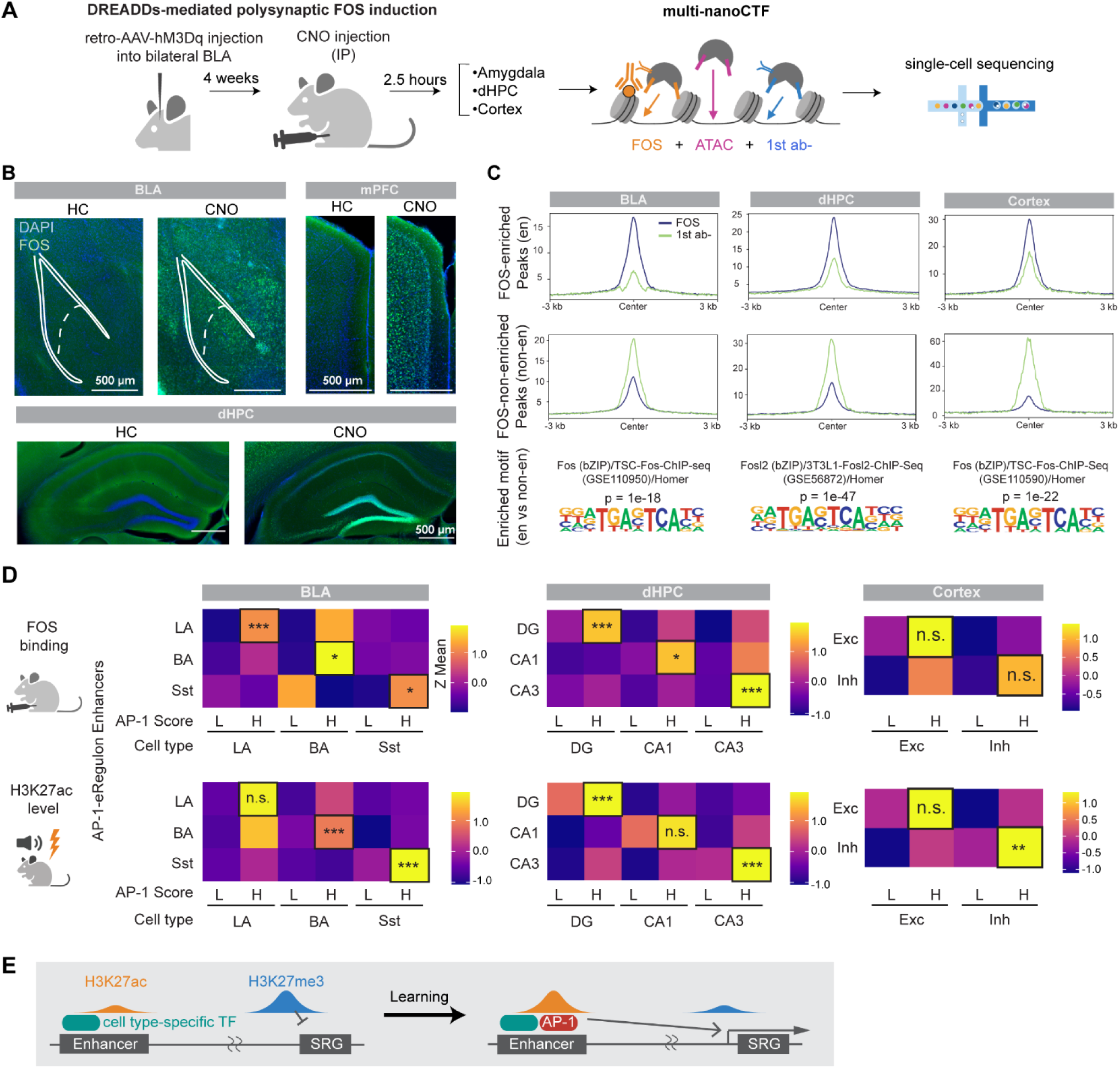
FOS multi-nanoCTF validates cell-type-specific AP-1 enhancer activation. (A) Scheme for multi-nanoCTF for FOS followed by polysynaptic FOS induction via excitatory DREADDs. Multi-nanoCTF jointly profiles chromatin accessibility, FOS occupancy, and matched first-antibody-minus background in the same nuclei using modality-barcoded nanobody–Tn5 reagents. (B) Representative immunofluorescence images showing systemic FOS induction 2.5 h after CNO-mediated activation of retrograde hM3Dq-expressing neurons in the BLA, dHPC, and mPFC. Green, FOS; blue, DAPI. Scale bars, 500 μm. (C) Top and middle: Aggregate plots showing FOS and 1st ab-minus signal in FOS-enriched or FOS-non-enriched ATAC-based peaks. Bottom: AP-1 (FOS) motifs were identified as representative motifs by comparing FOS-enriched peaks vs FOS-non-enriched peaks. (D) Heatmaps showing normalized FOS occupancy (top; multi-nanoCTF after DREADDs) and H3K27ac levels (bottom; multi-nanoCT after tone aversive conditioning) at cell-type-specific AP-1 eRegulon enhancers. Rows represent enhancer sets assigned to each cell-type-specific AP-1 eRegulon, and columns represent neurons of each cell type stratified into AP-1 motif score-low (L) and score-high (H) states. Cell-type-specific AP-1 eRegulon enhancers showed preferential FOS enrichment and H3K27ac accumulation in the matched AP-1-high cell types. FOS occupancy was quantified as background-corrected FOS signal relative to the first-antibody-minus control. Because broad DREADD-induced polysynaptic FOS expression did not produce uniform AP-1-associated regulatory activity, neurons were stratified by AP-1 motif score into AP-1-low (L) and AP-1-high (H) regulatory states. Statistical significance was assessed using a one-sided stratified permutation test with 5,000 permutations, comparing the observed matched-cell-type enrichment with randomly sampled enhancer sets while preserving enhancer set size and overall detection-frequency bins. (E) Model of SRG regulation upon learning in neuronal ensembles. At a basal state, distal enhancers are primed by cell type-specific transcription factors (TFs), and gene-proximal regions are covered by Polycomb-repressive H3K27me3. Upon learning, enhancers are activated by AP-1 binding accompanied with H3K27ac accumulation, leading to Polycomb remodelling and SRG induction.

To generate a high-confidence FOS-occupancy validation dataset with sufficient signal for single-cell analysis, we used a hM3Dq-based chemogenetic strategy to drive distributed neuronal activation ^40,41^. Because the BLA serves as a highly connected structure capable of extensive polysynaptic activation, we bilaterally injected a retrograde AAV-hM3Dq into the BLA to target its broad afferent network ^42,43^. Following Clozapine N-Oxide (CNO) administration, we triggered robust, trans-synaptic network FOS expression, whereas control mice showed only baseline-level FOS expression. This approach leveraged multi-nanoCTF profiling (Figure 4B).

We identified candidate FOS-bound regulatory regions by classifying scATAC-seq peaks based on FOS signal relative to the 1st antibody-minus background (Figure 4C). Despite the inherent sparsity of single-cell binding profile, FOS-enriched peaks were strongly enriched for AP-1 motifs, indicating that this strategy captured *bona fide* FOS-bound regions. Consistently, genome browser tracks showed reproducible FOS enrichment above background signals across brain regions, indicating a favourable signal-to-background ratio (Figure S5D). Furthermore, at the single-cell level, AP-1 motif scores defined by ATAC-seq modality were positively associated with FOS signals, validating increased FOS occupancy in nuclei with high AP-1-associated chromatin accessibility (Figure S5E).

We next examined FOS occupancy at cell-type-specific AP-1 eRegulon-associated enhancers identified above. These enhancers showed preferential FOS enrichment in their matched neuronal cell types compared with other cell types (Figure 4D, top). This enrichment reached statistical significance in the BLA and dHPC, whereas the mPFC showed the same directional trend with a smaller effect size.

We next asked whether FOS-bound AP-1 eRegulon enhancers acquire an active enhancer state upon learning. To this end, we used the matched H3K27ac modality from multi-nanoCT datasets of tone fear conditioned animals and compared Cond High with Cond Low neurons within each corresponding cell type. This showed that these AP-1-bound enhancers were also preferentially marked by increased H3K27ac in Cond High cells of the corresponding cell types (Figure 4D, bottom).

Together, these results suggested a distal enhancer regulatory layer that is distinct from gene-proximal Polycomb-H3K27me3 remodelling, in which cell-type-specific AP-1 eRegulon enhancers coordinate chromatin accessibility, FOS occupancy, H3K27ac enrichment, and target-gene induction (Figure 4E).

### Identification of learning-associated genes (LAGs)

Having defined activity-associated transcriptional and epigenomic programs in ChromTRAP-defined neuronal ensembles, we next asked which components were selectively associated with associative learning rather than baseline neuronal activity. We therefore sought to define learning-associated genes (LAGs) as AP-1-regulated SRGs specifically induced in Cond High neurons compared with HC High neurons.

To identify candidate AP-1-regulated SRGs, we integrated multimodal epigenetic and transcriptional regulatory layers. We first intersected ChromTRAP-defined DEGs with AP-1 eRegulon-associated genes, focusing on excitatory neurons because inhibitory neurons yielded too few DEGs for robust downstream analysis (Figure 5A and S6A). This identified 55 AP-1-regulated SRGs in the BLA, 128 in the dHPC, and 18 in the mPFC. We then compared these genes between Cond High (i.e., putative engram formed by learning) and HC High neurons to define LAGs as the subset preferentially induced by conditioning rather than by non-specific home-cage activity (Figures 5B and S6B; Table S6). This yielded 26 LAGs in the BLA, 49 in the dHPC, and 5 in the mPFC. LAGs included plasticity-related genes such as *Sorcs3*, *Pcdh15*, and *Bdnf*, and transcription-associated regulators including *Mapk4* and *Smad3*, supporting their relevance to memory-associated transcription. *Scg2,* recently implicated in memory-related plasticity, was the only LAG shared across all three regions (Figure 5C and S6C)^32^.

**Figure 5.**
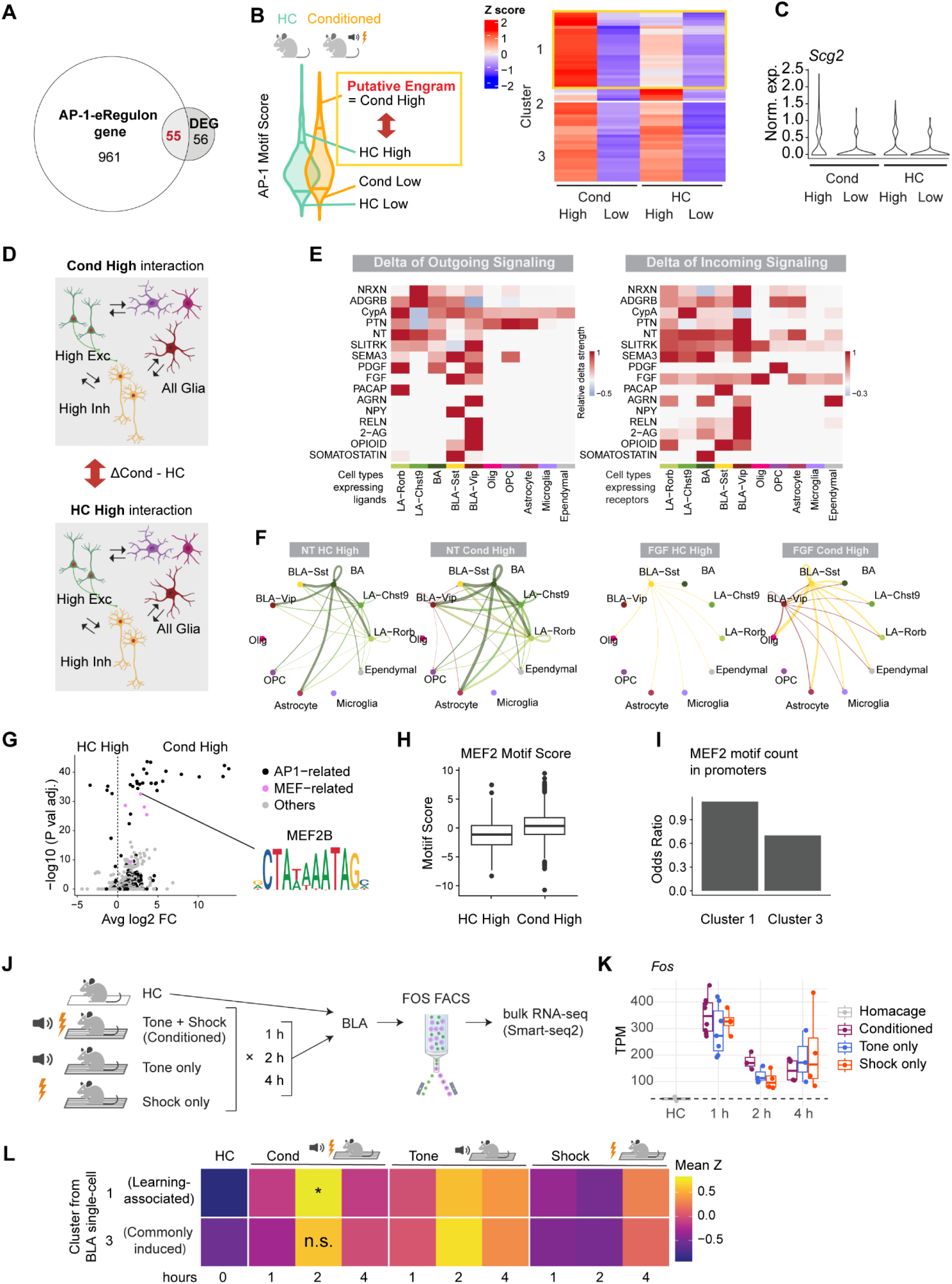
Characterization of learning-associated genes (LAGs). (A) Venn diagram showing the overlap between AP-1-eRegulon-associated genes and ChromTRAP-defined DEGs in BLA excitatory neurons. Overlapping genes were defined as AP-1-regulated SRGs. (B) Left: Scheme for identifying learning-associated genes (LAGs). AP-1-regulated SRGs preferentially expressed in Cond High (i.e., putative engram cells formed by learning) compared to home-cage (HC) High cells are defined as LAGs. Right: Heatmap showing expression levels of AP-1-regulated SRGs across ChromTRAP-defined BLA excitatory neurons. Cluster 1 genes were defined as LAGs by their preferential expression in Cond High neurons, whereas Cluster 3 genes were classified as commonly induced genes shared between Cond High and HC High neurons. (C) Violin plot showing *Scg2* expression in ChromTRAP-defined BLA excitatory neurons. (D) Scheme for intercellular molecular interaction analysis combining ChromTRAP and CellChat. CellChat-inferred interactions between Cond High and HC High populations, while retaining glial populations as potential interaction partners. (E) Heatmaps showing changes in outgoing (top) and incoming (bottom) signalling strength between Cond High and HC High populations. Pathways detected in either Cond High or HC High are shown. Overall stronger signalling in Cond High neurons suggests learning-associated enhancement of intercellular molecular interactions. Pathway abbreviations are listed in Table S7. (F) Circle plots showing inferred cell-type-specific ligand–receptor interactions for neurotrophin (NT) and growth-factor (FGF) signalling. Edge thickness indicates inferred signalling strength. Excitatory and inhibitory neurons are shown in green- and yellow-toned colours. (G) Volcano plot showing differential transcription factor motif scores (i.e., chromVAR scores) between Cond High and HC High neurons. MEF2-related motifs are shown in pink, AP-1-related motifs in black, and other motifs in grey. (H) Box plot showing MEF2 motif scores (i.e., chromVAR score) in Cond High and HC High neurons in BLA excitatory neurons. (I) Bar plot showing odds ratio of MEF2 motif occurrence in promoters of Clusters 1 and 3 genes from Figure 5B, representing LAGs and commonly induced genes. (J) Experimental scheme for bulk RNA-seq validation of LAGs. Mice were exposed to tone conditioning, tone-only, or shock-only paradigms and sacrificed 1, 2, or 4 h after. BLA NeuN+ nuclei from HC mice and FOS+/NeuN+ nuclei from stimulated mice were isolated by FACS and subjected to bulk RNA-seq. (K) Box plot showing *Fos* expression across paradigms and time points. (L) Heatmap showing expression of LAGs and commonly induced genes, defined by ChromTRAP-based single-cell multiomics analysis (see Figure 5B), in the bulk FOS-FACS RNA-seq validation dataset. Statistical significance was assessed using the self-contained gene set test fry (limma), comparing the conditioning 2 h group with the other 2 h conditions. *P < 0.05, fry gene set test.

For downstream epigenetic analyses, we prioritized BLA excitatory neurons because they showed the clearest distinction between Cond High and HC High neurons and provided sufficient LAG resolution for chromatin-level characterization (Figures 5B and S6B).

Moreover, the BLA is a canonical circuit node for aversive tone conditioning ^44,45^, providing a biologically well-defined setting to examine how learning-associated transcription is coupled to transcriptional and epigenetic regulation.

We found that more than half of these LAGs overlapped Polycomb-associated H3K27me3 domains, linking this learning-associated subset to the gene-proximal Polycomb regulatory layer identified above (Figure S6D).

Together, the comparison between ChromTRAP-defined Cond High and HC High neurons separates conditioning-biased transcriptional programs from transcription associated with baseline activity.

### Characterization of LAG programs

We further investigated LAG programs. First, the enrichment of cell–cell interaction-related genes in ChromTRAP-defined neuronal ensembles (Figure 2H) prompted us to examine whether activated neurons exhibit altered intercellular signaling states. To address this, we integrated ChromTRAP with CellChat ^46,47^, a bioinformatic tool that infers intercellular molecular interactions by mapping ligand-receptor gene expression patterns from single-cell transcriptomes. We compared CellChat-inferred signalling pathways between Cond High and HC High populations, and found that signalling strength was generally higher in Cond High neurons (Figures 5D, 5E, S6E, and S6F), indicating learning-associated enhancement of molecular interactions. Furthermore, outgoing secreted-ligand pathways showed cell-type specificity, with neurotrophin and FGF signalling predominantly derived from excitatory and inhibitory neurons, respectively, whereas recipient populations were more diverse (Figure 5E and 5F).

Next, to identify candidate regulators that distinguish LAGs from commonly induced activity genes (i.e., genes also induced in HC High neurons), we integrated ChromTRAP with transcription factor motif screening (Methods). We examined motifs preferentially associated with Cond High against HC High neurons, and identified enrichment of MEF2 family motifs (Figure 5G, 5H, and S6G). Using FIMO-based motif counting ^48^, we found that MEF2 motifs were more strongly enriched in LAG promoters than in promoters of commonly induced genes (Figure 5I). *Mef2c* showed the highest basal expression, identifying it as a putative regulator of the LAG program (Figure S6H).

Together, these results indicate that Cond High neurons engage a learning-associated gene regulatory program characterized by enhanced intercellular ligand–receptor communication and MEF2-associated transcriptional regulation.

### LAGs mark learning-specific transcriptional programs

We next tested whether LAGs identified by single-cell multiomics analysis are preferentially induced by aversive tone conditioning, rather than by tone or shock exposure alone. We performed bulk RNA-seq on FOS/NeuN-positive neurons isolated from the BLA by FACS at 1, 2, and 4 h after tone-only, shock-only, or tone-conditioning with home-cage mice as controls (Figure 5J).

In conditioned animals, the proportion of FOS-positive cells increased at 1 h after conditioning (about 4 %), declined by 2 h, and returned to near home-cage levels by 4 h (Figure 2B). Consistently, *Fos* mRNA levels in sorted neurons peaked at 1 h and declined thereafter, yet it still remained detectable with bulk mRNA-seq at 4 h (Figure 5K).

Differential expression analysis relative to home-cage controls identified 79, 159, and 62 upregulated genes at 1, 2, and 4h after conditioning, respectively (Figure S6I; Table S8). Hierarchical clustering revealed an early general activity-response cluster enriched for IEGs and induced across all three paradigms (cluster 1 in Figure S6J). Conversely, later clusters (clusters 2 and 3) contained genes preferentially induced by tone conditioning compared with tone-only or shock-only stimulation at 2-4 h (Figure S6J). These conditioning-biased genes significantly overlapped with single-cell-defined LAGs (Figure S6K).

As a complementary validation, we quantified single-cell-defined LAG expression in the bulk FOS-FACS RNA-seq dataset (Figure 5L). While bulk RNA-seq lacks neuronal subtype resolution, it provides an orthogonal validation whether LAGs are preferentially induced by conditioning. Because FOS-positive cell recovery was markedly reduced at 4 h, we focused on the 2 h time point, where both cell recovery and transcriptional responses were robust. LAGs exhibited significantly higher expression following tone conditioning compared to tone-only or shock-only conditions. This conditioning-biased induction was not observed for genes commonly induced in Cond High and HC High neurons.

Together, these complementary single-cell and bulk analyses support LAGs as a conditioning-biased AP-1-associated SRG program, preferentially engaged by tone–shock association rather than by baseline activity or sensory/aversive stimulation alone during early memory formation. This framework links LAG induction to chromatin organization, enhanced intercellular signaling, and MEF2-family regulatory input, and provides a mechanistic basis for how neuronal ensembles engage learning-associated transcription.

## Discussion

Memory formation involves a transition from IEG induction to SRG activation in activated neuronal ensembles, yet this phase has remained difficult to access, because FOS-like activity markers decay rapidly, whereas genetic reporters (e.g., TRAP) require several hours for detectable expression. We addressed this gap by developing ChromTRAP, an scATAC-seq-based strategy that uses AP-1 motif accessibility as a retrospective trace of recent neuronal activation. When applied 4 h after conditioning, ChromTRAP enriched for recently activated, memory-associated neuronal ensembles across multiple brain regions (Figure 2A, right). Although ChromTRAP does not establish functional engram identity, it enabled ensemble-defined integration of multimodal regulatory layers. By combining scMultiome, multi-nanoCT, and the newly developed multi-nanoCTF platforms, we resolved multilayered epigenetic and transcriptional programs within the same neuronal ensembles (Figure 1A). We proposed a proximal–distal epigenomic logic for activity-dependent transcription in neuronal ensembles: H3K27me3 remodeling at SRG proximal regions and AP-1-bound H3K27ac-marked distal enhancers jointly shape cell-type-specific transcriptional programs (Figure 4E).

Our analyses also link learning-associated programs to the concept of “pre-configuration” of memory engram cells ^4,49^. Previous studies have shown that memory-related neuronal recruitment (memory allocation) is biased toward neurons with higher excitability, including CREB-dependent excitability, connectivity and chromatin plasticity ^4,50–52^. The comparison between ChromTRAP-defined Cond High and HC High neurons provides an entry point for understanding how such activity-associated states are transformed into learning-associated programs. Cond High neurons were characterized by the induction of LAGs, which are conditioning-biased subset of AP-1-targeted SRGs. These neurons also showed enhanced intercellular molecular interactions, and enrichment of MEF2-family motifs (Figure 5). MEF2 factors, particularly MEF2C, are linked to human cognition through rare loss-of-function neurodevelopmental disorders^53^ and resilience against age-related cognitive decline, including Alzheimer’s disease^54,55^. Mechanistically, MEF2 regulates activity-dependent transcription, synaptic refinement/pruning, and suppression of excessive potentiation ^56–60^. At the behavioral level, *Mef2c* deletion impairs learning and memory despite increased excitatory synapse number and transmission, whereas acute MEF2 gain- and loss-of-function bidirectionally modulates spatial and fear memory ^58,59^. Collectively, MEF2 may help balance activity-induced plasticity and contribute to the specificity of memory-associated neuronal ensembles. This possibility suggests that learning-associated transcription may shape engram not only by enhancing selected connections but also by homeostatically refining the broader activity-induced response.

Beyond providing access to the SRG expression window, ChromTRAP may provide a broader advantage of chromatin-based readouts for identifying rare and transient cell states in single-cell datasets. scRNA-seq-based detection of recently activated neurons can be limited by technical and biological sparsity: although *Fos* and other activity-responsive gene transcripts are informative markers, their detection in single-cell transcriptomics can be affected by stochastic dropout, while memory engrams represent only a small fraction of neurons. Together, these factors can reduce sensitivity for identifying activated cells and limit the downstream differential analysis. scATAC-seq-based ChromTRAP addresses this challenge by reading a distributed chromatin footprint of transcription factor activity rather than relying on a small set of transcripts. Because FOS/AP-1 acts across thousands of activity-regulated enhancers, AP-1 motif accessibility can be aggregated into a continuous per-cell activity score. This robustly identified recently activated neurons across protocols, brain regions, and neuronal types. Thus, ChromTRAP shows how single-cell chromatin profiling can complement, and in some cases surpass, transcriptome-only readouts for detecting rare cell states. A previous work showed that these AP-1 footprints are substantially reduced by 24 h after neuronal activation but can partially persist, suggesting that such chromatin traces may remain informative over this timescale of SRG induction ^7^. More broadly, this strategy may be useful for dissecting experience-dependent neuroplasticity beyond memory, including auditory maternal behaviour, sensory-dependent development and damage responses, as well as other rare transcription factor-driven cell states in non-neuronal cells across biological contexts ^60–63^.

These findings also define several limitations and future directions. First, we characterized putative engram cells during learning, but the functional consequences of their states during later recall remain unresolved. In particular, it will be important to determine how AP-1-bound enhancers and Polycomb-remodelling influence the specificity and composition of recall-induced transcriptional responses. Combining ChromTRAP with longitudinal activity tagging (e.g., genetic TRAP, CalLight, FLiCRE) ^5,10,14–19,64,65^, recall-induced IEG mapping, or calcium imaging could bridge this gap. Second, while CellChat nominated specific signaling pathways (NT, IGF, FGF), resolving their spatial organization will require spatial omics-based approaches which may be integrated with ChromTRAP analysis. Third, although this study focused on AP-1, neuronal activity also induces other IEG-encoded transcription factors such as EGR and NPAS4. The correlation between AP-1, EGR- and NPAS4-related motif activities (Figure S2C) suggests that AP-1-based ChromTRAP captures a broad activity-dependent regulatory state. Extending multi-nanoCTF to these factors could reveal how distinct IEG-driven programs diversify neuronal ensembles. Lastly, LAG characterization of this study focused on a relatively simple tone aversive conditioning paradigm. Future applications to paradigms with distinct cognitive and circuit demands, such as trace conditioning, could test whether the identity and regional specificity of LAG programs change when cortical and hippocampal mechanisms for attention and temporal association are more strongly engaged.

Together, these findings establish ChromTRAP, integrated with single-cell multiomics, as a technical and conceptual framework for linking experience to chromatin state and gene regulation in memory-associated neuronal ensembles, with future potential for dissecting diverse forms of experience-dependent plasticity.

### Limitations of ChromTRAP

ChromTRAP has methodological constraints. First, it requires the integration of a single-cell chromatin accessibility modality introducing greater experimental and computational complexity than transcriptome-only single-cell approaches. Second, ChromTRAP is designed to identify and molecularly characterize neuronal ensembles, but unlike genetic activity-tagging methods (e.g.,TRAP2), it does not provide transgene-based access for subsequent manipulation of the same cells. Finally, consistent with other IEG-based approaches, ChromTRAP should not be interpreted as a direct measure of neuronal firing. Rather, AP-1-centered chromatin traces are best viewed as markers of activity-linked regulatory and plasticity-associated states, and should ideally be integrated with complementary physiological, longitudinal, or functional readouts.

## RESOURCE AVAILABILITY

### Lead contact

Further information and requests for resources and reagents should be directed to and will be fulfilled by the lead contact, Taro Kitazawa.

### Materials availability

All unique reagents generated in this study are available from the lead contact without restriction. Reagents, chemicals, animals, and software used in this study are available in the Methods.

### Data and code availability

Sequencing raw data and processed data and codes are deposited at ArrayExpress and GitHub.

## ACKNOWLEDGEMENT

We thank the members of the Kitazawa laboratory, especially Y. K. Yumiko, M. Vestergaard, and S. W. Andersen for their kind support of experiments and analysis. We thank G. Castelo-Branco (Karolinska Institute), ME. Greenberg (Harvard Medical School), T. Kim (Aarhus University), S. Nabavi (Aarhus University), J. Nagai (RIKEN), K. Hagihara (National Institutes of Natural Sciences), S. Josselyn, and Paul Frankland (SickKids) for critical discussion regarding the project. We acknowledge the Danish Single-Cell Examination Platform (CellX), especially L. Lin (Aarhus University), for providing access to 10x Controller. CellX was established with support from the Danish Research Agency through the Danish national research infrastructure program (5229-0009B). We appreciate that cell sorting was performed at the FACS Core Facility (Aarhus University). Next-generation sequencing was performed by the MOMA Core Center, Department of Molecular Medicine, Aarhus University Hospital, Denmark or BGI Genomics, China. The authors acknowledge the Bioimaging Core Facility, Health, Aarhus University, Denmark, for the use of equipment and support. We used BioRender for figure preparation. We used ChatGPT to help coding and English proofreading. This study was supported by European Research Council ERC-StG MemoPlasticGenomics 101039734, Lundbeckfonden DANDRITE R361-2020-2654, Lundbeckfonden R412-2022-639, Novo Nordisk Foundation 0096135, Novo Nordisk Foundation 0088740, Aarhus Universitets Forskningsfond AUFF-E-2024-9-40, Japan Science and Technology Agency (JST) PRESTO JPMJPR24N6, and Danish National Research Foundation PROMEMO DNRF133.

## AUTHOR CONTRIBUTIONS

Conceptualization: KI, VK, TK. Methodology: KI, VK, IF, TK. Investigation: KI, VK, IF, TK. Visualization: KI, VK. Funding acquisition: TK. Project administration: TK. Supervision: TK. Writing – original draft: KI, VK, IF. Writing – review & editing: TK

## Materials and Methods

### Mice

#### Animals

Male mice of the strain C57BL/6JRj were purchased from Janvier Labs, France. Mice were purchased at 6–8 weeks of age. All mice were housed in a 12 hour light/dark cycle at 23°C and had ad libitum food and water access. Mice were housed 4 per cage. All procedures that involved the use of mice were approved by the Danish Animal Experiment Inspectorate (permit numbers: 2020-15-0201-00421 and 2023-15-0201-01431). All experiments were performed using adult male mice. Male mice were used in this study to establish and benchmark ChromTRAP and associated single-cell multiomic workflows under a controlled experimental setting, while avoiding additional sources of variation related to sex chromosome composition, X-chromosome dosage or inactivation, and sex-dependent transcriptional and chromatin differences that could complicate integrated genomic analyses.

### Aversive tone conditioning

All behavioural testing was performed between 1 PM and 6 PM. The apparatus consisted of an open-top cage (24 × 20 × 30 cm) with metal floor bars placed inside a soundproof cubicle (55 × 60 × 57 cm) (Ugo Basile, Italy). Mice were conditioned in Context A using two pairings of a 25 s, 7 kHz sine-wave tone (conditioned stimulus, CS) that co-terminated with a 1.5 s, 0.7 mA footshock (unconditioned stimulus, US). Separate cohorts of mice were used to assess short-term memory (STM) and long-term memory (LTM) recall in a novel context (Context B) at 4 h and 24 h after conditioning, respectively. After 2 min of acclimation to the new context, mice were presented with two CS presentations without footshock. The intertrial interval ranged from 90 to 120 s during both conditioning and testing sessions.

Context B was a modified version of Context A. The stainless-steel bars were covered with a white plastic insert, and a handful of Alpha-dri (Innovive) was placed on top. The chamber was scented using cotton embedded with a few drops of isoamyl acetate (Sigma-Aldrich, W205508).

An additional group was exposed to the CS without any US presentation on day 1 (tone-only group) and was re-exposed to the same CS in Context B on the following day, as described above. Behavioural responses were recorded with a top-view camera, and freezing was scored automatically using ANY-maze software (version 7.2; Stoelting Europe, Ireland). Freezing percentage was defined as the time spent freezing during the CS presentation divided by the CS duration and multiplied by 100. During conditioning, freezing percentage was calculated for each CS presentation. During recall, freezing percentage was calculated as the average across the two CS presentations. Pre-CS freezing percentage was calculated as the average freezing time during the 25 s preceding each CS presentation.

For sequencing experiments (smart-seq2 and scMultiome and multi-nanoCT), mice in the conditioning group were conditioned as described above, except that after conditioning they were kept in isolation for 4 h before sacrifice and brain collection for sequencing. Two control groups were included for smart-seq2 experiments: one group was exposed twice to the CS (tone only), and the other was exposed only to the US (shock only). Both control groups were also isolated for 4 h before sacrifice, and brains were harvested for sequencing experiments.

### Statistics for behavioral analysis

Statistical analyses were performed by using GraphPad Prism 10. All the data are represented as mean ± SEM, and they were tested for normality using Shapiro–Wilk and D’Agostino–Pearson normality test. If the data represented a normal distribution, a parametric test was used. The statistical methods and the corresponding p-values are reported in the figure legends.

### Viral Delivery and Chemogenetic Activation

Prior to surgery, mice received a subcutaneous injection of buprenorphine (Temgesic, 0.3 mg/mL; 0.1 mg/kg) for analgesia. Anesthesia was induced and maintained using isoflurane (IsoFlo vet 100%, Zoetis). Standard stereotaxic surgical procedures were employed to expose the skull and perform a craniotomy.

Mice received bilateral injections of AAV-retro/2-mCaMKIIα-hM3D(q) (titer: 6 × 10¹² vg/mL, diluted 1:1 in PBS) targeting the basolateral amygdala (BLA). An injection volume of 0.2 to 0.5 µL per hemisphere was delivered at the following stereotaxic coordinates relative to Bregma: anteroposterior (AP) –1.6 mm, mediolateral (ML) ±3.45 mm, and dorsoventral (DV) −3.5 to −4.1 mm from the skull surface.

For immunostaining experiments, water-soluble clozapine N-oxide dihydrochloride (CNO; Hello Bio, Cat. No. HB6149) was administered three weeks after AAV delivery by intraperitoneal injection (2 mg/kg in a 400 µL volume), and mice were sacrificed 2 h after CNO administration. For multi-nanoCTF experiments, CNO was administered four weeks after AAV delivery using the same dose and route, and mice were sacrificed 2.5 h after CNO administration.

### Microdissection of BLA, dorsal hippocampus, mPFC, and SSC

Mice were deeply anesthetized with isoflurane, decapitated, and brains were placed in a stainless-steel brain matrix on ice to cut 1 mm sections along the anterior-posterior axis.

Sections were transferred into ice-cold Slicing buffer (26 mM NaHCO3, 2.5 mM KCl, 1.25 mM NaH2PO4, 10 mM MgSO4, CaCl2 0.5 mM, Glucose 11 mM, Sucrose 234 mM) bubbled with 5 % CO2 and 95 % O2, and BLA, dorsal hippocampus, mPFC, and somatosensory cortex (SSC) were separated under a dissection microscope. Tissue was snap-frozen in liquid nitrogen and stored at −80 °C.

### Nuclear extraction

Frozen tissues were transferred to a pre-chilled Dounce homogenizer and immediately lysed in ice-cold buffer. For scMultiome and Smart-seq2 experiments, NP-40 nuclei extraction buffer was used, containing 20 mM Tris-HCl pH 7.6, 25 mM KCl, 5 mM MgCl₂, 0.25 M sucrose, 1 mM DTT, and RNase inhibitor. RNase inhibitor (Thermo Fisher Scientific, Cat. No. EO0384) was added at 1 U/µL for scMultiome experiments and 0.4 U/µL for Smart-seq2 experiments. For multi-nanoCT experiments, antibody buffer was used, containing 20 mM HEPES pH 7.5, 150 mM NaCl, 2 mM EDTA, 0.01% digitonin, 0.01% NP-40, 0.5 mM spermidine, 1% BSA, and protease inhibitor (Roche, Cat. No. 11836170001). The buffer volume was adjusted according to tissue size of the tissues: BLA, 500 µL; mPFC, 500 µL; hippocampus, 1,000 µL. On ice, the tissue was immediately homogenized 5 times with the loose homogenizer, then 10 times with the tight one. Nuclei were filtered (miltenyibiotec, 130 041 407) and were transferred into a new tube.

### Single-cell Multiome

#### Nuclear cleaning by OptiPrep

For scMultiome using whole nuclei, nuclei were cleaned by density-gradient centrifugation with OptiPrep (OptiPrep, Sigma D1556). Each biological sample consisted of pooled material: two mice for HC and three for the Tone Conditioning 4 hours. N = 2 per condition. To the extracted nuclei, 1.5X volume of ice-cold G50 Buffer was added and mixed gently by inverting tubes. 1-2 mL (depending on the size of tissues) of G30 Buffer was slowly underlaid. Nuclei were centrifuged for 30 min at 1000g at 4°C. Floating debris was carefully removed, and nuclei were resuspended in 500 µL ice-cold 1% BSA Buffer. Nuclei were centrifuged for 5 min at 500g at 4°C, and resuspended in 500 µL ice-cold 1% BSA Buffer. Nuclei were washed once more. For permeabilization, we added 0.1X Lysis Buffer and slowly mixed it by pipetting 5 times. After 1 min incubation on ice, ice-cold 500 µL 1% BSA Buffer was added and slowly mixed it by pipetting 5 times. Nuclei were filtered (30 um) to a new tube, took 10 µL for cell counting. Nuclei were centrifuged for 5 min at 500g at 4°C. After removal of supernatant, ice-cold 1X Diluted Nuclei Buffer (included in 10x Multiome kit) was added to make a dilution of over 3k nuclei/ul. Nuclei were counted using a hemocytometer and the volume to use for the next step was decided.

<u>1X Tris Buffer 1</u>: 20mM Tris pH 7.6, 25 mM KCl, 5 mM MgCl2, 0.25M Sucrose

<u>6X Tris Buffer-1</u>: 120 mM Tris pH7.6, 150 mM KCl, 30 mM MgCl2

<u>Tris Buffer-2:</u> 11 mM Tris pH7.6, 11 mM NaCl, 3.3 mM MgCl2

<u>G50 Buffer</u>: 1X 6X Tris Buffer-1, 50% OptiPrep, 1mM DTT, 1U/uL RNAse inhibitor

<u>G30 Buffer:</u> 0.4X 1X Tris Buffer-1, 0.6X 6Xtris Buffer-1, 30% OptiPrep, 1mM DTT, 1U / µL RNAse inhibitor

<u>1% BSA Buffer</u>: 1X Tris Buffer-2, 1% BSA, 1 mM DTT, 1U / µL RNAse inhibitor

<u>0.1X Lysis Buffer</u>: 1X Tris Buffer-2, 1% BSA, 1 mM DTT, 1U/ µL RNAse inhibitor, 0.001% Digitonin, 0.01% NP40, 0.01% Tween20

### Immuno-FACS-sorting for FOS or 7AAD+singlet

For FOS-FACS sorting Multiome analysis, four biological samples were pooled for one biological replicate. N = 1. For 7AAD+singlet FACS sorting (mPFC Cond 4h replicate 1), three biological samples were pooled to one biological replicate. N=1.

Extracted nuclei were centrifuged for 5 min at 500g at 4°C in a swing bucket. Pellet was resuspended in 500 µL 1% BSA PBS. After 5 min incubation on ice, pipetting mix.

Antibodies (anti-FOS: ab208942 conjugated with Alexa 488, 1/1,000, anti-NeuN: ab190565 conjugated with Alexa 647) were added, and rotated at 4°C for 1 h. Nuclei were centrifuged for 5 min at 500g at 4°C in a swing bucket, and resuspended in 1% BSA PBS with 1 µg/ml 7AAD after 5 min incubation on ice. Just before sorting, nuclei were filtered into 5 mL tube (Corning, 352235). FACS was performed with Bigfoot (Thermofisher). NeuN+/FOS+ or 7AAD+singlet nuclei were sorted into Nuclei Buffer (10x Genomics, 2000207) supplemented with 1 mM DTT, 1 U/ul RNase inhibitor (Sigma, 3335399001), and 1% BSA. After FACS, nuclei were centrifuged for 5 min at 500g at 4°C in a swing bucket. Supernatant was removed leaving 5-10 µL and 5 µL was used for downstream processing.

### Library preparation and sequencing

The DNA and RNA libraries were prepared according to the manufacturer’s protocol. 10x Multiome ATAC–seq and RNA–seq libraries were sequenced in paired-end mode (ATAC: R1–i7–i5–R2 = 60–8–24–60; RNA: R1–i7–i5–R2 = 27–10–10–90) on an Illumina NovaSeq X Plus platform (Illumina).

### Basic analysis for whole nuclei sequencing data

Raw data were processed using *CellRanger ARC* (v2.0.2, 10x Genomics) with default settings. Different samples and replicates in the same brain regions were aggregated using *CellRanger ARC aggr* (Hippocampus and BLA: default settings; mPFC: “norm = none”).

Low-quality cells were removed using *Seurat* ^66^ v5.3.0 and *Signac* ^67^ v1.15.0 according to the following quality-control criteria. Across all analyzed regions, cells were retained if they had 500–8,000 RNA features, TSS enrichment greater than 1, and nucleosome signal less than 1.5. Region-specific ATAC count thresholds were applied as follows: 300–100,000 counts for BLA, 500–100,000 counts for mPFC, and 400–100,000 counts for hippocampus.

RNA counts were normalized using *SCTransform*^68^ with default settings. Cell clustering was performed in three rounds. First, cells were separated into three major groups: excitatory neurons, inhibitory neurons, and glial cells. Cell types were identified based on the expression of marker genes: *Slc17a7* and *Slc17a6* for excitatory neurons; *Gad2* for GABAergic neurons; and *Slc1a3, Mal, Mog, Pdgfra, C1qb,* and *Sostdc1* for glial cells.

Second, doublets were removed, defined as cells simultaneously expressing marker genes from different lineages (e.g., *Gad2* and *Slc17a7*). Third, subclustering of each major group was performed. At each step, clustering was conducted using *RunPCA, FindNeighbors, FindClusters*, and *RunUMAP* (*Seurat*).

After each round, raw counts of the selected cells were obtained, and *SCTransform* and peak calling with *MACS2* were re-run. For mPFC, before *SCTransform*, the Seurat object was split for each sample. Before clustering, the split objects were merged by *IntegrateLayers* (option: “method = HarmonyIntegration”). For PFMs, JASPAR2024.sqlite3 ^69^ was used.

ChromVAR ^27^ score was calculated with *RunChromVAR* function (*Seurat*) (option: “genome = BSgenome.Mmusculus.UCSC.mm10”).

### Basic analysis for FACS-sorted FOS+ nuclei sequencing data

Raw data was processed using *CellRanger ARC*. Ambient RNA was normalized using *SoupX* ^70^ v1.6.2 before building Seurat object. Cells were retained if they had 300–8,000 RNA features, TSS enrichment greater than 1, and 500–100,000 ATAC counts. Following peak calling using *MACS2* and *SCTransform*, clustering was performed as described in the previous section. For hippocampus, batch effect was normalized using *Harmony*.

### Motif enrichment analysis comparing conditioning versus HC

Differentially accessible (DA) peaks were calculated with *FindMarkers* function (*Seurat*) using “peak” assay by specifying conditioning sample as ident.1 and HC sample as ident.2 (parameters: test.use = “wilcox”). The top 1000 p-value peaks were identified as DA peaks. *FindMotifs* function (*Seurat*) was used to search for enriched motifs within DA peaks.

### Identification of DEGs using whole nuclei sequence data

Neurons in the top 10% and bottom 10% of AP-1 motif scores in each condition were used for this analysis. For BLA and hippocampus, differentially expressed genes were identified using *FindMarkers* in *Seurat* with the Wilcoxon rank-sum test and min.pct = 0.01. Genes with an absolute average log2 fold change of at least 1 and an adjusted P value less than 0.01 were considered differentially expressed. For mPFC, differential expression analysis was performed using *edgeR* ^71^ v4.4.2 after summing raw count values across cells within each ChromTRAP-defined group and replicate. Low-expression genes were filtered with *filterByExpr*, library sizes were normalized with *calcNormFactors*, and differential expression was assessed with a quasi-likelihood F-test using *glmQLFit* and *glmQLFTest*. Genes with false discovery rate (FDR)< 0.05 were considered significantly differentially expressed and were classified as up- or down-regulated according to the sign of the log2 fold change.

DEGs were identified for three comparisons: Cond High versus Cond Low, Cond High versus HC Low, and HC High versus HC Low. Genes significantly up-regulated in any of these comparisons in the same cell types were used as ChromTRAP-associated DEGs for downstream analyses, including DEG counting, gene ontology analysis, Polycomb analysis, and overlap analysis with AP-1-eRegulon genes.

### Identification of DEGs to benchmark ChromTRAP

Neurons below the 10th percentile in the HC group were defined as HC Low, and neurons above the 90th percentile in the Cond 4 h group were defined as Cond High. Pseudobulk raw counts were then calculated for each condition using *AggregateExpression* function in *Seurat*. For the FACS-sort versus HC Low comparison, mitochondrial and ribosomal genes were excluded from the analysis to minimize the effect of ambient RNA. Raw counts were normalized using the *DESeq* function in *DESeq2*, and DEGs were identified using *results* function in *DESeq2*. Genes with a log2 fold change greater than 1 and an adjusted p value less than 0.07 were defined as DEGs.

### Gene Ontology (GO) analysis

Up-regulated DEGs in excitatory neurons or inhibitory neurons in BLA, mPFC, and dorsal hippocampus were aggregated and subjected to GO analysis with R package *clusterProfiler* ^72^ v4.14.6 using following parameters: OrgDb = "org.Mm.eg.db", pvalueCutoff = 0.05, qvalueCutoff = 0.05, ont = "MF".

### Gene Regulatory Network Analysis with Scenic+

Gene regulatory network analysis was performed using *SCENIC+* ^35^. For each cell type, cells in the top 20th percentile and bottom 20th percentile of the AP1 score in conditioned samples were identified as “Cond High” and “Cond Low” and were used for analysis.

Consensus peaks and topics were identified using *pycisTopic* v2.0a0. Topics enriched in Cond High neurons, together with differentially accessible regions identified from imputed accessibility profiles by comparing Cond High and Cond Low neurons, were used for *SCENIC+* v1.0a2 analysis. Enhancer–gene relationships were then inferred using *SCENIC+*. Among the resulting eRegulons, those containing AP-1-associated motif annotations were defined as AP-1 regulons. Cell types which did not contain AP-1 regulons were excluded from the analysis. In the mPFC, because enhancer features were similar between PV and Sst cells, AP-1 regulons from these two cell types were integrated. eRegulons were visualized using Cytoscape ^73^ v3.10.4. Cell-type-specific enhancers were defined as those for which the mean accessibility in Cond High neurons of the annotated cell type was at least 0.1 higher than that in the other Cond High neurons in other cell types, while the mean accessibility in Cond Low cells of the same annotated cell type was lower than the 70th percentile within that annotation group. Signal intensities of scATAC-seq within AP-1-eRegulon enhancers were quantified using the *FeatureMatrix* function in *Signac*. For visualization of ATAC and RNA levels, top 20^th^ percentile and bottom 20th percentile were used.

### Identification of Cell Type-specific Motifs and counting within AP-1-eRegulon enhancers

The *FindMarkers* function was used by specifying “chromVAR” slot with default settings. Top 50 motifs were selected as candidate for cell-type-specific AP-1-eRegulon enhancer regulator. Motif occurrences were counted using *FIMO* in *meme* ^48^.

### Identification of Learning-Associated Genes (LAGs)

AP-1-regulated SRGs were defined as genes overlapping between DEGs and AP-1 eRegulon genes across cell types. To identify learning-associated genes, mean expression levels of AP-1-regulated SRGs were calculated across four groups: Cond High, Cond Low, HC High, and HC Low, using the data layer of the SCT assay. Expression values were row-wise Z-scored and subjected to hierarchical clustering. Distances were calculated using the *dist* function in the R stats package v4.4.2 with Euclidean distance, and clustering was performed using the *hclust* function with the Ward.D2 method.

### CellChat

Neurons in the top 10% of AP-1 motif scores in each condition were defined as “High” and used for this analysis. Cell–cell molecular interactions were inferred using the R package *CellChat* ^47^ v2.2.0 according to the developer’s tutorials (Inference and analysis of cell–cell communication using CellChat). Separate CellChat objects were generated for each based on SCT normalized single-cell RNA-seq expression matrices. Merged CellChat objects were constructed to enable pairwise comparison of signaling networks between conditions using the *mergeCellCha*t and *rankNet* functions. Differentially enriched pathways were identified as those with significantly higher signaling strength scores in the conditioned group compared to controls (p < 0.05). Network centrality scores (incoming and outgoing strengths) were computed for each cell type using netAnalysis_computeCentrality.

To visualize relative changes in pathway activity across cell types, we computed the difference of normalized centrality matrices between groups and plotted the results as heatmaps using *ComplexHeatmap* ^74^ v2.22.0. Color gradients were scaled per pathway and capped at the 2–98 % quantile to highlight the relative change in signaling strength.

The Network strength of each ligand-receptor pair was plotted with *netVisual_aggregate* function (options: layout = "circle").

### Differential motif activity analysis comparing Cond High versus HC High

Excitatory neurons in the top 20% of AP-1 motif scores within each condition were defined as Cond High and HC High and used for this analysis across brain regions. Differential motif activity between Cond High and HC High neurons was assessed using FindMarkers in Seurat with the chromVAR assay. Cond4h cells were compared against HC cells using logistic regression with nCount_peaks included as a latent variable. Motifs with an average log2 fold change of at least 1 and an adjusted P value less than 0.001 were classified as up-regulated, whereas motifs with an average log2 fold change of −1 or less and an adjusted P value less than 0.001 were classified as down-regulated. All other motifs were classified as unchanged.

To highlight selected motif patterns, JASPAR CORE motif metadata and visual inspection of motif logos were used to annotate MADS box factor motifs as MEF-related motifs and bZIP or AP-1-like motifs as AP-1-like motifs. MEF-related motifs were highlighted in violet, AP-1-like motifs in black, and all other motifs in gray.

### MEF2 motif count in LAG promoters

Gene annotation data (Mus musculus GRCm38.102) were obtained from Ensembl in GTF format and processed using R packages *rtracklayer* v1.66.0. Transcript entries were extracted, and transcription start sites (TSSs) were specified as the most upstream coordinate for transcripts on the positive strand and the most downstream coordinate for those on the negative strand. Promoter regions were defined as −2 kb to +1.5 kb relative to each TSS, taking strand orientation into account, and gene body regions were assigned accordingly. Then, promoters associated with SRGs were extracted using gene_name columns in the GTF file. *FIMO* was used to count occurrences of the MEF2 motif, which was obtained from JASPAR2024 (MA0660.1).

### Oligonucleotides

All oligonucleotides used in this study, including adapters, indexed adapters, sequencing primers, reverse-transcription primers, and template-switching oligonucleotides, are listed in Table S9.

### Multiplexed nanobody-mediated single-cell CUT&Tag

#### Biological Samples

For multi-nanoCT experiments targeting transcription factors (FOS, 1st ab-minus, and ATAC), tissues from chemogenetically activated mice were used. For dHippo rep1, two biological samples were pooled. In all other cases, each biological sample represented one biological replicate. For the cortex dataset, rep1 was derived from the mPFC, whereas rep2 was derived from the posterior somatosensory cortex (SSC).

### ATAC reaction

Extracted nuclei were centrifuged at 300 g for 3 min at 4°C. The pellet was resuspended in Antibody Buffer, using 550 µL for histone-modification samples and 1 mL for transcription factor samples. For permeabilization, nuclei were rotated at 4°C for 1 h and then centrifuged at 300 × g for 3 min. For histone-modification samples, nuclei were washed twice with 150 µL Wash Buffer (10 mM Tris, 10 mM NaCl, 5 mM MgCl₂, 1% BSA) before proceeding to the next step.

For both sample types, the pellet was resuspended in 150 µL ATAC Reaction Buffer containing 20 mM Tris pH 7.6, 10 mM MgCl₂, 20% dimethylformamide, 0.33× PBS, 0.01% Tween-20, and 0.01% digitonin, supplemented with 15 µL ME-A Tn5 transposase. Nuclei were incubated at 37°C for 30 min with shaking at 300 rpm. The reaction was stopped by adding 7.5 µL of 500 mM EDTA. Nuclei were then pelleted by centrifugation at 300 × g for 3 min at 4°C.

### Antibody Treatment and Tn5 Labeling

The nuclei were resuspended in 300 µL Antibody Buffer. Primary antibodies were added and incubated overnight at 4°C with rotation. The following primary antibodies were used: anti-H3K27ac (Abcam, ab6002), anti-H3K27me3 (Abcam, ab4729), and anti-FOS (Santa Cruz Biotechnology, sc-253). No-primary-antibody controls were processed in parallel. After centrifugation at 300 × g for 2 min at 4°C, the pellet was resuspended in 300 mM NaCl Buffer supplemented with both mouse and rabbit nanobody–Tn5. Transcription factor samples received 225 µL buffer containing 13.5 µL of each nanobody–Tn5, whereas histone-modification samples received 150 µL buffer containing 9 µL of each nanobody–Tn5. The 300 mM NaCl Buffer consisted of 20 mM HEPES pH 7.5, 300 mM NaCl, 0.01% digitonin, 0.01% NP-40, 0.5 mM spermidine, 1% BSA, and protease inhibitor. Nuclei were rotated at room temperature for 1 h, washed with ice-cold 300 mM NaCl Buffer, and centrifuged as above. Sample-to-barcode assignments for each multiplexed nanoCT library are listed in Table S10.

Tagmentation was initiated by adding Tagmentation Buffer, consisting of 300 mM NaCl Buffer supplemented with MgCl₂ to a final concentration of 10 mM. Histone-modification samples received 100 µL Tagmentation Buffer, whereas transcription factor samples received 150 µL. Nuclei were mixed by pipetting and incubated at 37°C for 1 h with shaking at 300 rpm. The reaction was stopped by adding an equal volume of ice-cold DNB Stop Buffer to each sample. DNB Stop Buffer consisted of Nuclei Buffer (10x Genomics), 1% BSA, and 25 mM EDTA.

### FACS and Sample Aggregation

Histone-modification samples were resuspended in 200 µL 1% BSA DNB Buffer and centrifuged at 200 × g for 3 min at 4°C. The pellet was then resuspended in 500 µL 1% BSA DNB Buffer containing 1 µg/mL 7-AAD, and singlet nuclei were sorted by FACS. For transcription factor samples, nuclei were resuspended in 300 µL Antibody Buffer and incubated with anti-FOS antibody (ab208942 conjugated with Alexa 488, 1/1,000) for 1 h at 4°C with rotation. Nuclei were then centrifuged at 300 × g for 3 min at 4°C, resuspended in 500 µL 1% BSA DNB Buffer containing 1 µg/mL 7-AAD, and FOS-positive singlet nuclei were sorted by FACS. To avoid aggregation after sorting, 2% BSA DNB Buffer was used because samples were diluted by sheath fluid during FACS. Sorted nuclei were pooled by combining equal numbers of nuclei from each sample. The pooled nuclei were centrifuged at 500 × g for 5 min at 4°C, and the supernatant was removed to achieve a target concentration of 2,000 × n nuclei/µL in 1% BSA DNB Buffer, where n indicates the number of pooled samples. Nuclei were counted again before downstream processing.

### 10X ATAC Library Preparation

For each 10X Genomics ATAC reaction, n × 16,000 nuclei, where n indicates the number of pooled samples, were resuspended in 8 μL 1% BSA DNB Buffer and 7 μL ATAC Buffer. GEM generation, barcoding, and linear amplification were performed according to the 10X Genomics protocol, except that Phusion High-Fidelity DNA Polymerase (Thermo Fisher Scientific, F530S) was used in place of the Barcoding Enzyme. A second tagmentation step was performed in Tris-TD Buffer (20mM Tris pH7.6, 10 mM MgCl2, and 20% dimethyl formamide) with 0.1–0.2 μL ME-B Tn5, followed by purification with Ampure beads (1.6X), ethanol washes, and elution in 40 μL EB buffer. The library PCR was performed following the manufacturer’s protocol, except that 13–15 PCR cycles were used. In addition, samples were incubated at 72°C for 5 min before amplification to allow complementary DNA strand synthesis at the ME-B Tn5 adaptor ends.

### Sequencing

Libraries were sequenced for paired-end (R1-i7-i5-R2 = 40-10-41-40) with Illumina NovaSeqX plus (Illumina).

### Basic data analysis for multi-nanoCT for histone modification

*Nanoscope* ^26^ was used for primary analysis including FASTQ file demultiplexing for each modality, running *CellRanger ATAC*, re-cell calling, re-peak calling using *MACS2* with “-broad” options. Within the merged broad peaks among samples, each modality signal was counted with *FeatureMatrix* function (*Seurat*). Cells with Top 1 % and bottom 1 % logUMI, were excluded from the analysis. UMAP connecting the same cell in different modalities was plotted using *plotConnectModal* function (*Nanoscope*). Plot showing genome browser view was generated with *CoveragePlot* function (*Signac*). ChromVAR ^27^ score was calculated as described above in scMultiome using H3K27ac modality.

### Basic data analysis for multi-nanoCT for FOS

*Nanoscope* was used only for modality-specific FASTQ demultiplexing, because *CellRanger ATAC* could not be applied directly due to the sparsity of these datasets. For ATAC modality, demultiplexed FASTQ was subjected to *CellRanger ATAC.* For TF and 1st ab-minus, demultiplexed FASTQ files were trimmed using *Trimmomatic* with ILLUMINACLIP:/path_to_adapters/NexteraPE-PE.fa:2:10:10:8:TRUE, MINLEN:30, LEADING:3, TRAILING:3, and SLIDINGWINDOW:4:15, and reads were aligned to the mm10 genome using *Bowtie2* ^75^ v2.5.1 with default settings.

To restrict FOS and 1st ab-minus libraries to high-quality nuclei, *CellRanger ATAC*-called barcodes from the matched ATAC libraries were used as the valid cell barcode set. Aligned BAM files for FOS and 1st ab-minus libraries were then filtered using custom Python scripts with *pysam* (https://github.com/pysam-developers/pysam) v0.22.1 to retain only reads associated with *CellRanger ATAC*-called barcodes. In addition, only properly paired reads with both mates mapped were retained using *SAMtools* ^76^ v1.6. Also, PCR duplicates were removed using *MarkDuplicates* in *Picard* ^77^ v2.18.29 with REMOVE_DUPLICATES=true.

The resulting BAM files were further filtered to retain reads with mapping quality of at least 30 using *SAMtools*. Filtered BAM files were converted to fragment files for downstream *Signac/Seurat* analysis using *Sinto*.

For visualization, filtered BAM files were converted to smoothed bedGraph tracks using custom Python scripts. Because the FOS and 1st ab-minus datasets were sparse, signal was smoothed by adding coverage within a 200 bp window around both ends of each properly paired fragment. BedGraph files were converted to bigWig files using *bedGraphToBigWig* ^78^ v4. Genome browser tracks were visualized using IGV v2.16.0.

### Cell type annotation

Cell-type-specific marker peaks were identified from single-nuclei Multiome scATAC-seq data using *Signac* and *Seurat*. After LSI-based dimensionality reduction, UMAP visualization, and clustering of the peaks assay, RNA-based cell-type annotations were transferred to the ATAC modality. Differentially accessible peaks for each cell type were identified using *FindMarkers* with the Wilcoxon rank-sum test and min.pct = 0.1. Marker-peak sets were selected using average log2 fold-change thresholds of >0.7 for BLA excitatory neurons and >1 for all other neuronal and glial populations.

Marker peaks derived from the Multiome dataset were overlapped with multi-nanoCT peak sets and used for cell-type annotation. H3K27ac peaks were used for histone-modification multi-nanoCT datasets, whereas ATAC peaks were used for FOS multi-nanoCT datasets. For each marker-peak set, normalized signal was calculated as the summed signal over overlapping peaks divided by the total peak signal in each nucleus. Marker-peak enrichment patterns were visualized on UMAPs and used to assign cell-type identities to clusters.

### Peak call for H3K27me3 modality and gene assignment

For H3K27me3 peak calling, cell-type-filtered BAM files containing excitatory and inhibitory neuronal nuclei were generated using *filterbarcodes* in *Sinto* v0.10.1 (https://github.com/timoast/sinto). Broad H3K27me3 peaks were called from the filtered BAM files using *epic2* ^79^ v0.0.54 with the mm10 genome and an FDR threshold of 0.05. After peak calling, peak regions were inspected using genome browser tracks. Because some broad H3K27me3 domains appeared fragmented for technical reasons, nearby peaks separated by gaps of up to 10 kb were merged using the *merge* function in *BEDTools* ^80^ with the -d 10000 option.

Merged H3K27me3 broad peaks longer than 10 kb were assigned to genes using overlap-based annotation. Gene annotations were obtained from the Mus musculus GRCm38.102 GTF file. For each gene, a gene-plus-promoter interval was defined by combining the gene body with a promoter region spanning ±2 kb around the transcription start site. H3K27me3 broad peaks were overlapped with these intervals using *GenomicRanges* ^81^ v1.58.0. Peaks were assigned to all overlapping genes.

### Identification of FOS-enriched peaks and Motif Enrichment Analysis

To identify FOS-enriched peaks, FOS and 1st ab-minus signals were quantified within matched ATAC-seq peaks using *multicov* in *BEDTools*. Peaks with a total count greater than 8 across FOS and first-antibody-negative control libraries were retained for analysis. Differential enrichment between FOS and 1st ab-minus signals was assessed using *edgeR*. Trimmed mean of M values (TMM) normalization factors were calculated using *calcNormFactors*. Because the 1st ab-minus library was expected to have lower overall signal by experimental design, equal library sizes were then assigned to the FOS and control libraries before model fitting. A negative binomial model was fitted using *glmFit* with a fixed dispersion corresponding to BCV = 0.3, followed by likelihood-ratio testing using *glmLRT*. Peaks with FDR < 0.05 were classified as FOS-enriched peaks, and all other tested peaks were classified as non-enriched.

Motif enrichment analysis was performed using *findMotifsGenome.pl* in *HOMER* ^82^ v5.1 with the mm10 genome and -size given. To identify motifs enriched in FOS-enriched peaks, non-enriched peaks were used as the background.

### Quantification of multi-nanoCT signals within arbitrary peak sets

To quantify multi-nanoCT signals within predefined genomic regions, peak or enhancer coordinates were converted to genomic ranges and used as input features for *FeatureMatrix* in *Signac*. Fragment counts for each modality, including ATAC, H3K27ac, H3K27me3, FOS, and 1st ab- control were calculated within the specified regions using the corresponding fragment files.

### Immunostaining

Immunostaining was performed as previously described ^83^, with the following modification: sections were incubated with anti-FOS antibody (clone 9F6; Cell Signaling Technology, Cat. No. 2250) at 1:1000 dilution, followed by Alexa Fluor 488 goat anti-rabbit secondary antibody (Invitrogen, A11008) at 1:1000 dilution. Images were acquired using an upright widefield slide scanner (VS120, Olympus). For visualization, stitching-related background intensity offsets in the green channel were corrected in Fiji ^84^ by subtracting the background intensity difference measured across tile boundaries.

### Smart-seq2

#### Immuno-FACS-sorting for FOS

Each biological replicate corresponded to one biological sample. Immuno-FACS sorting was performed as described above, except that sorted nuclei were collected directly into Lysis Buffer from the Single Cell RNA Purification Kit (Norgen, #51800).

### RNA purification and library preparation

RNA was purified using the Single Cell RNA Purification Kit (Norgen Biotek, #51800) according to the manufacturer’s protocol. During RNA purification, on-column DNase treatment was performed using the RNase-Free DNase I Kit (Norgen Biotek, #25710).

RNA-seq libraries were prepared using a Smart-seq2-based protocol followed by Nextera XT library preparation. Briefly, up to 1 ng RNA was used for reverse transcription. RNA was mixed with dNTPs and oligo-dT30VN primer (sequence; Table S9), incubated at 72°C for 3 min, and immediately placed on ice. Reverse transcription was performed using SuperScript II reverse transcriptase (Invitrogen, 18064022), RNase inhibitor, betaine, MgCl₂, DTT, and template-switching oligo. The reverse-transcription reaction was run at 42°C for 90 min, followed by 10 cycles of 50°C for 2 min and 42°C for 2 min, and a final incubation at 72°C for 15 min.

Pre-amplification of cDNA was performed using KAPA HiFi HotStart ReadyMix (Roche Diagnostics, KK2602) and ISPCR primer. PCR was performed at 98°C for 3 min, followed by 13–18 cycles of 98°C for 20 s, 67°C for 15 s, and 72°C for 6 min, with a final extension at 72°C for 5 min. The number of PCR cycles was adjusted according to the input cell number. Half of the pre-amplification reaction was purified using AMPure beads at a 0.8× ratio and eluted in 10 mM Tris pH 7.6. cDNA concentration was measured using Qubit, and fragment size distribution was assessed using a Bioanalyzer for selected samples.

For library preparation, pre-amplified cDNA was diluted to 0.16 ng/µL, and 1.25 µL of diluted cDNA was used for tagmentation with the Nextera XT DNA Library Preparation Kit (Illumina, FC-131-1024). Tagmentation was performed by adding 1.25 µL Amplicon Tagment Mix and 2.5 µL Tagment DNA Buffer, followed by incubation at 55°C for 10 min. The reaction was stopped by adding 2.5 µL Neutralize Tagment Buffer and incubating at room temperature for 5 min.

Library PCR was performed by adding 3.75 µL Nextera PCR Master Mix and 2.5 µL IDT for Illumina DNA/RNA UD indexes (Illumina, 20027213). PCR was performed at 72°C for 3 min and 95°C for 30 s, followed by 15–20 cycles of 95°C for 10 s, 55°C for 30 s, and 72°C for 60 s, with a final extension at 72°C for 5 min. The number of PCR cycles was adjusted according to the input cell number. Libraries were purified using AMPure beads at a 0.6× ratio and eluted in 12 µL of 10 mM Tris pH 7.6. Library concentration was measured using Qubit, and library size distribution was assessed using a Bioanalyzer for selected samples. Libraries were sequenced in paired-end mode with DNBSEQ (BGI).

### Data analysis

Adaptors and low-quality bases were trimmed from paired-end RNA-seq reads using Trimmomatic *v0.39*^85^ with ILLUMINACLIP:NexteraPE-PE.fa:2:10:10, LEADING:3, TRAILING:3, SLIDINGWINDOW:4:15, and MINLEN:30. Paired reads passing filtering were used for downstream analysis. Trimmed reads were aligned to the Mus musculus GRCm38 primary assembly using *STAR* v 2.7.10b ^86^. Only uniquely mapped reads were retained by setting --outFilterMultimapNmax 1. Aligned reads were sorted using *SAMtools*. Gene-level read counts were generated using featureCounts from *subread* v2.0.1 ^87^ with the Mus musculus GRCm38.102 gene annotation. Fragments were counted using the -p option, and reads overlapping multiple features were assigned using the -O option. Numbers of reads retained at each processing step are provided in Table S11.

Genes were considered lowly expressed and excluded from the analysis if their maximum raw count across all samples was 1 or less.Transcripts per million (TPM) values were calculated from the filtered raw count matrix and used for visualization. Differentially expressed genes between each conditioned time point and home-cage controls were identified using *DESeq2* v1.64.0 ^88^. Genes with an absolute log2 fold change greater than 1.5 and an adjusted P value less than 0.0001 were defined as differentially expressed genes. For clustering, mean TPM values were calculated across biological replicates. Genes differentially expressed in the conditioned groups were row-wise Z-scored and hierarchically clustered using Euclidean distance and Ward.D2 linkage. Enrichment of overlap between bulk RNA-seq LAGs and single-cell Multiome LAGs was assessed using a one-sided Fisher’s exact test. The background universe was defined as the intersection of genes retained after bulk RNA-seq filtering and genes included in the single-cell Multiome GRN and FindMarkers input gene sets, resulting in 20,460 genes.

---

**Supplementary Figure 1.**
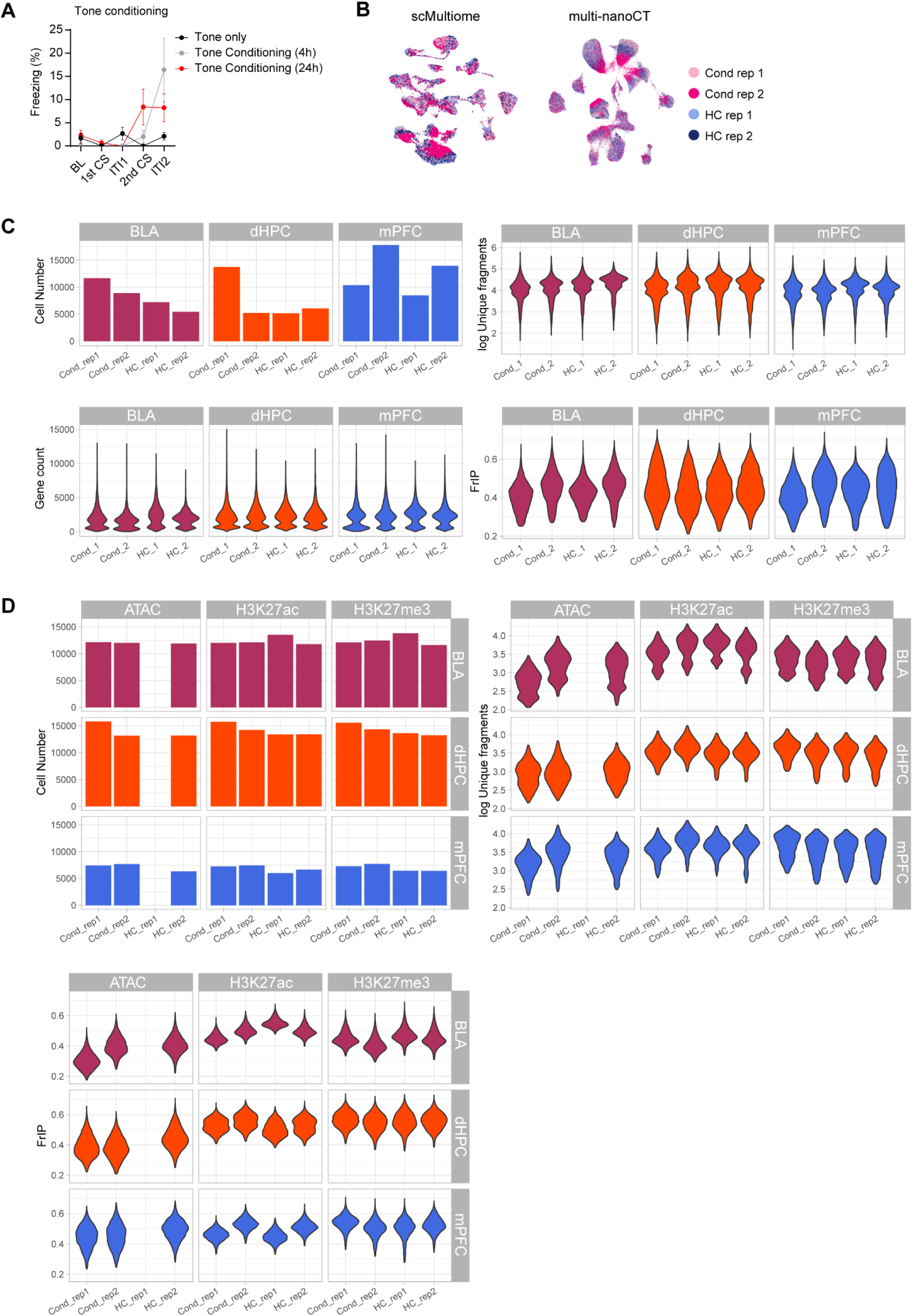
Behavioral validation and quality control of single-cell multi-omics datasets. (A) Freezing during tone-conditioning sessions in tone-only mice and mice analyzed 4 h or 24 h after tone conditioning. Mice were exposed to the tone in a different context. BL, baseline; CS, conditioned stimulus; ITI, inter-trial interval. (B) UMAP representing each replicate in scMultiome and multi-nanoCT data. (C, D), Quality-control metrics for scMultiome (C) and multi-nanoCT (D).

**Supplementary Figure 2.**
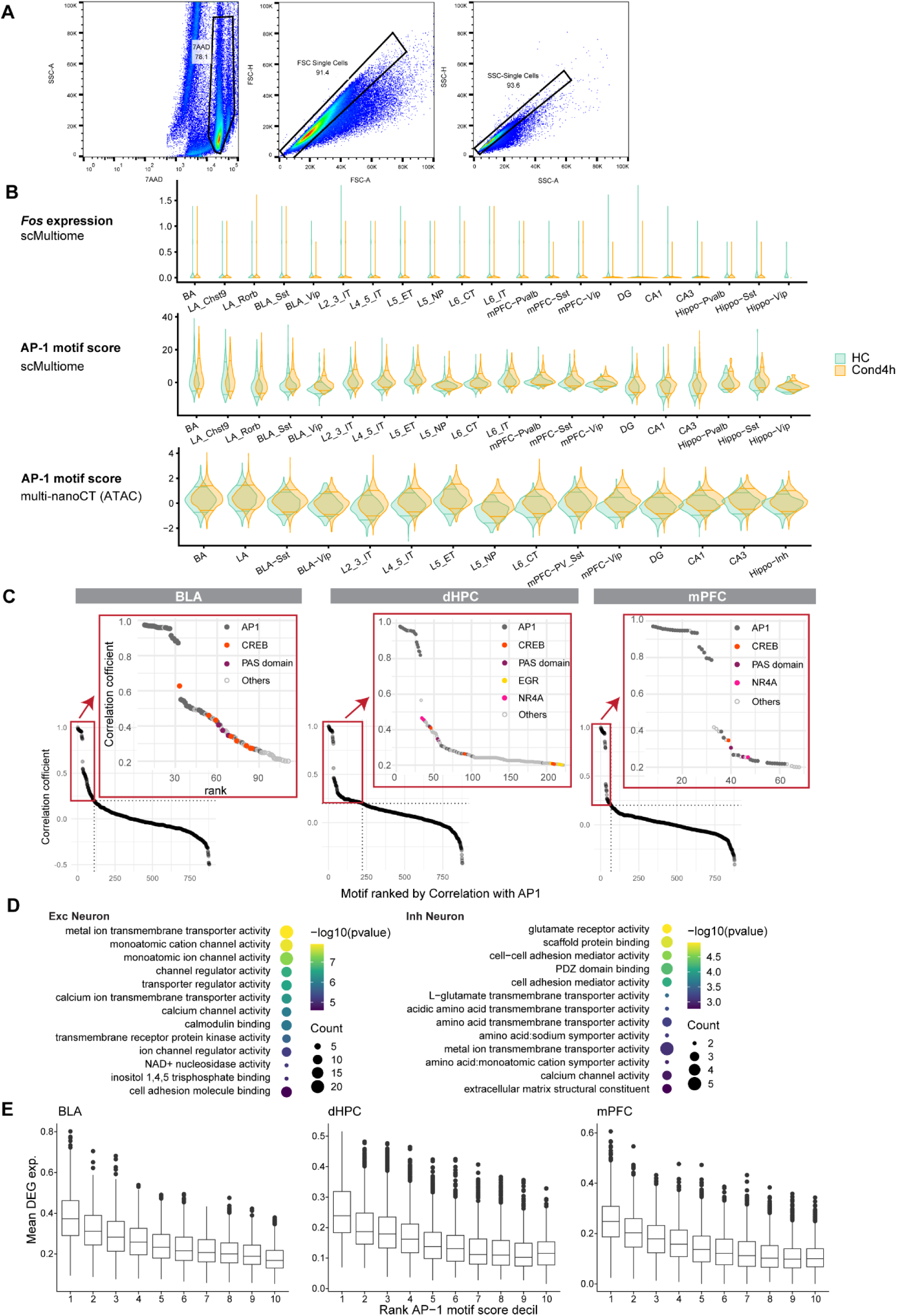
AP-1 motif accessibility supports ChromTRAP-based activity detection. (A) Representative FACS gating strategy for isolation of nuclei. 7-AAD-positive nuclei were selected, followed by forward scatter (FSC)- and side scatter (SSC)-based singlet gates. (B) *Fos* mRNA expression in scMultiome data (*top*), AP-1 motif scores in scMultiome data (*middle*), and AP-1 motif scores in multi-nanoCT data (*bottom*) across neuronal subtypes and brain regions. The horizontal lines indicate the *1*0th and 90th percentiles. The upper 90th percentiles was shifted upwords in Cond 4h. (C) Ranked plots of Pearson’s correlations between chromVAR scores for each transcription factor motif and the AP-1 motif score calculated using MA1141.2 in Cond High excitatory neurons. Insets show the top positively correlated motifs, indicating positive correlations between AP-1 and other well-known activity-associated transcription factor families (e.g., CREB, PAS, EGR, NR4A). (D) GO terms enriched in downregulated genes after aggregation across brain regions for excitatory and inhibitory neurons. (E) Box plots showing mean expression of ChromTRAP-defined DEGs across AP-1 motif score deciles in BLA, dorsal hippocampus and mPFC. Mean DEG expression was highest in cells with the top AP-1 motif scores, whereas intermediate score cells showed only modest increases as compared to lowest score cells. This indicates that top-scoring neurons retain a clear transcriptional contrast even when intermediate-score neurons, rather than HC Low or Cond Low neurons, are used as the baseline.

**Supplementary Figure 3.**
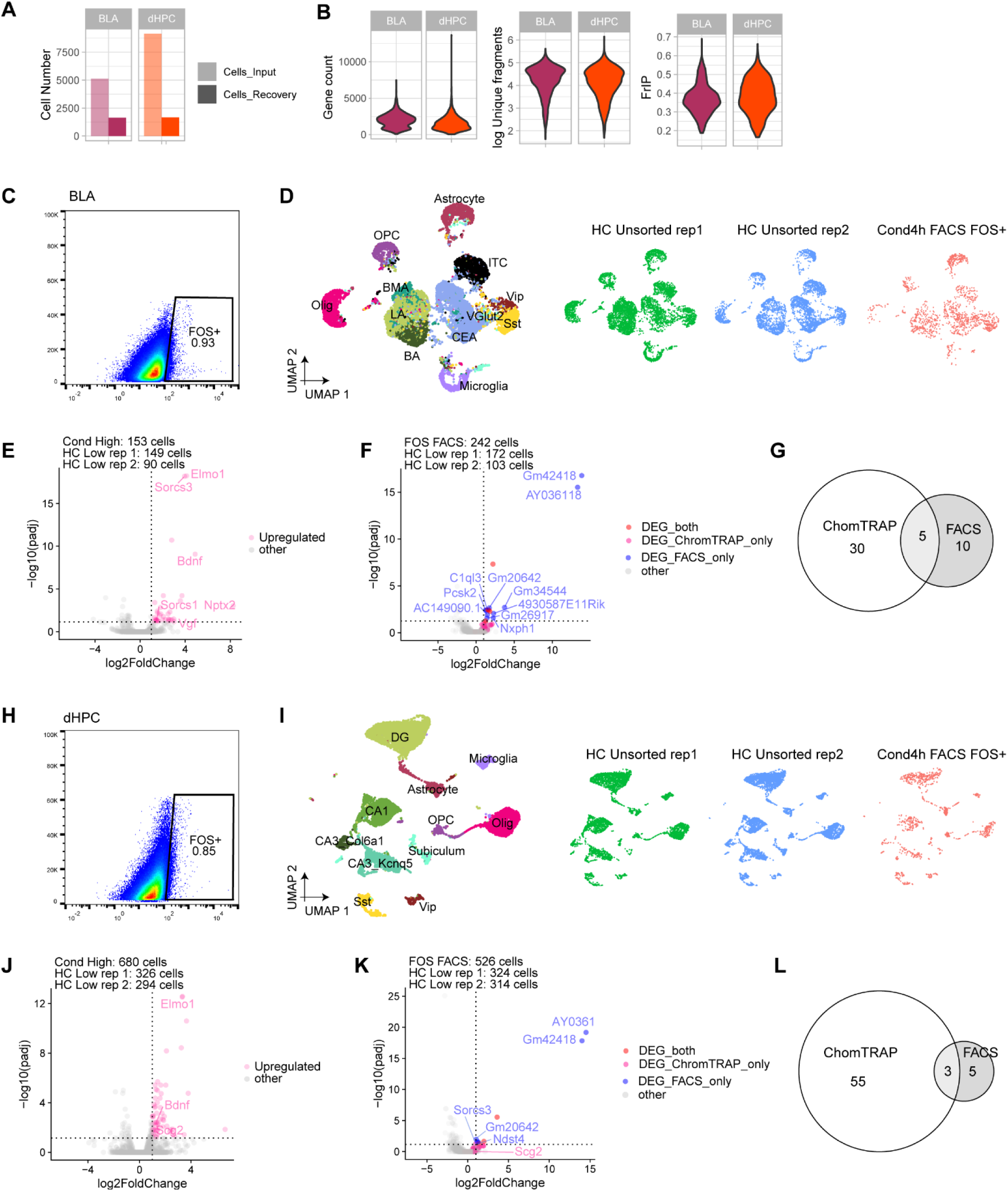
FOS-FACS Multiome validates ChromTRAP-defined differentially expressed genes. (A) Number of cells loaded and Cell Ranger-estimated recovered cells. (B) Quality control metrics for FOS-FACS Multiome datasets, including loaded and recovered cell numbers, unique fragments and FRiP scores. (C, H) Representative FACS plots showing FOS+ nuclei sorting from BLA (C) and dHPC (H) 4 h after conditioning. (D, I) UMAPs showing co-embedding of FOS-FACS Multiome nuclei with the corresponding unsorted scMultiome reference datasets in BLA (D) and dHPC (I) used for ChromTRAP analysis in this study. (E, J) Volcano plots showing DEGs identified by ChromTRAP-defined Cond High compared to HC Low in BLA (E) and dHPC (J). (F, K) Volcano plots comparing FOS-FACS-isolated neurons with HC Low neurons in BLA (F) and dHPC (K). DEGs shared with Cond High neurons are shown in red, Cond High-only DEGs in pink, and FOS-FACS-only DEGs in blue. Although ChromTRAP-defined Cond High DEGs tended to show positive fold changes in the FOS-FACS dataset, they rarely reached statistical significance, indicating limited sensitivity of FOS-FACS for detecting this activity-associated program. Furthermore, significant DEGs from FOS-FACS were enriched for poorly characterized Gm/Rik genes. (G, L) Venn diagrams showing overlap between ChromTRAP- and FOS-FACS-derived DEGs.

**Supplementary Figure 4.**
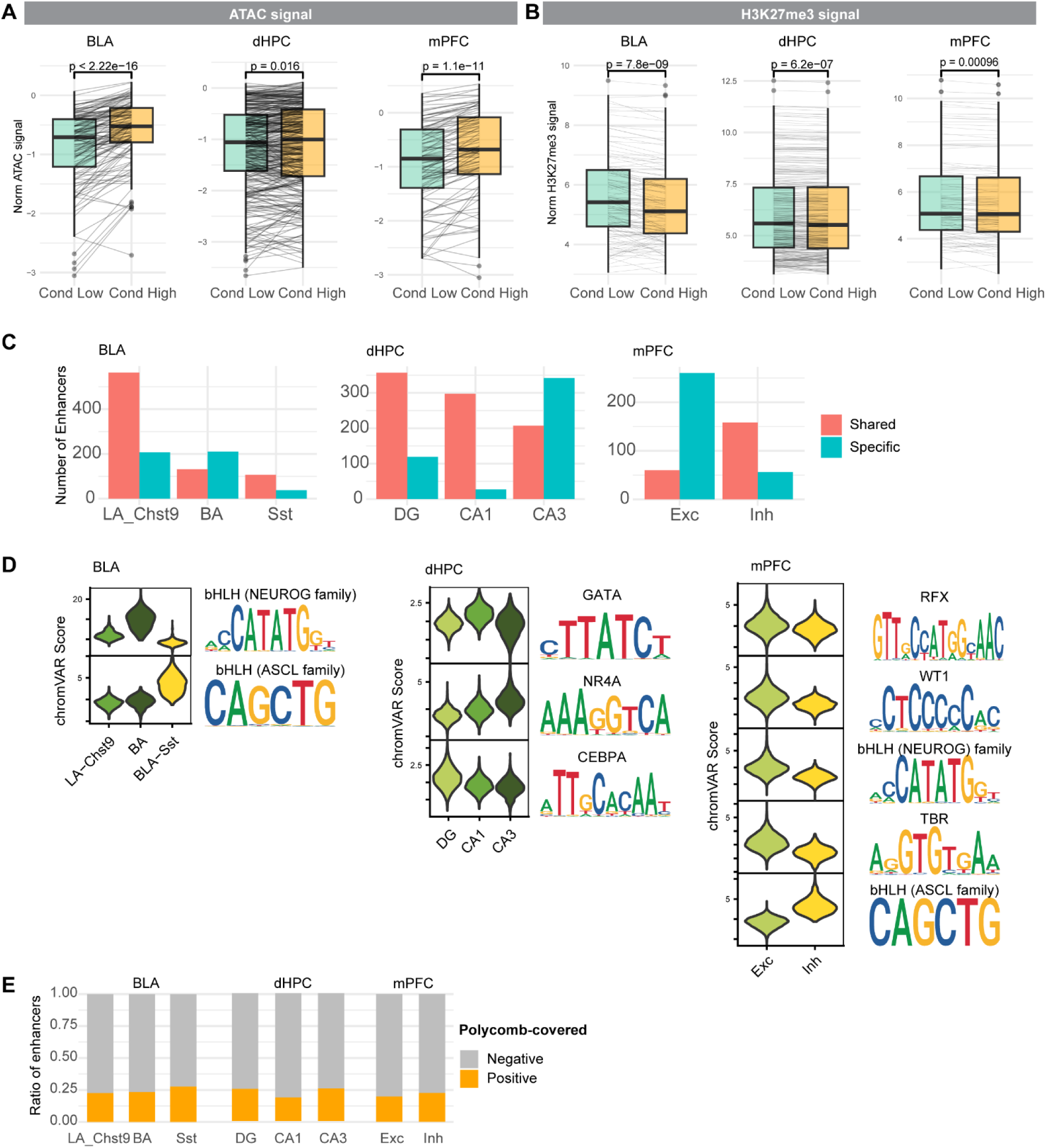
Epigenetic regulation of activity-response genes. (A, B) Box plots showing ATAC (A) and Polycomb-H3K27me3 (B) signals within Polycomb-H3K27me3-covered activity-dependent DEGs in Cond Low and Cond High neurons across brain regions. Lines connect the matched genes between conditions. Excitatory neurons are shown. Statistical significance was assessed by paired Wilcoxon signed-rank test. (C) Bar plots showing the number of shared and cell-type-specific AP-1 eRegulon enhancers across neuronal cell types and brain regions. Enhancers were classified as cell-type-specific when they showed preferential accessibility gain in the matched cell type (see Methods). (D) Violin plots showing chromVAR motif scores for cell-type-enriched transcription factor motifs identified from basal accessibility differences among neuronal cell types. Sequence logos indicate the corresponding enriched motifs. (E) Bar plots showing the fraction of AP-1 eRegulon enhancers overlapping Polycomb-H3K27me3-covered genomic regions across neuronal cell types and brain regions.

**Supplementary Figure 5.**
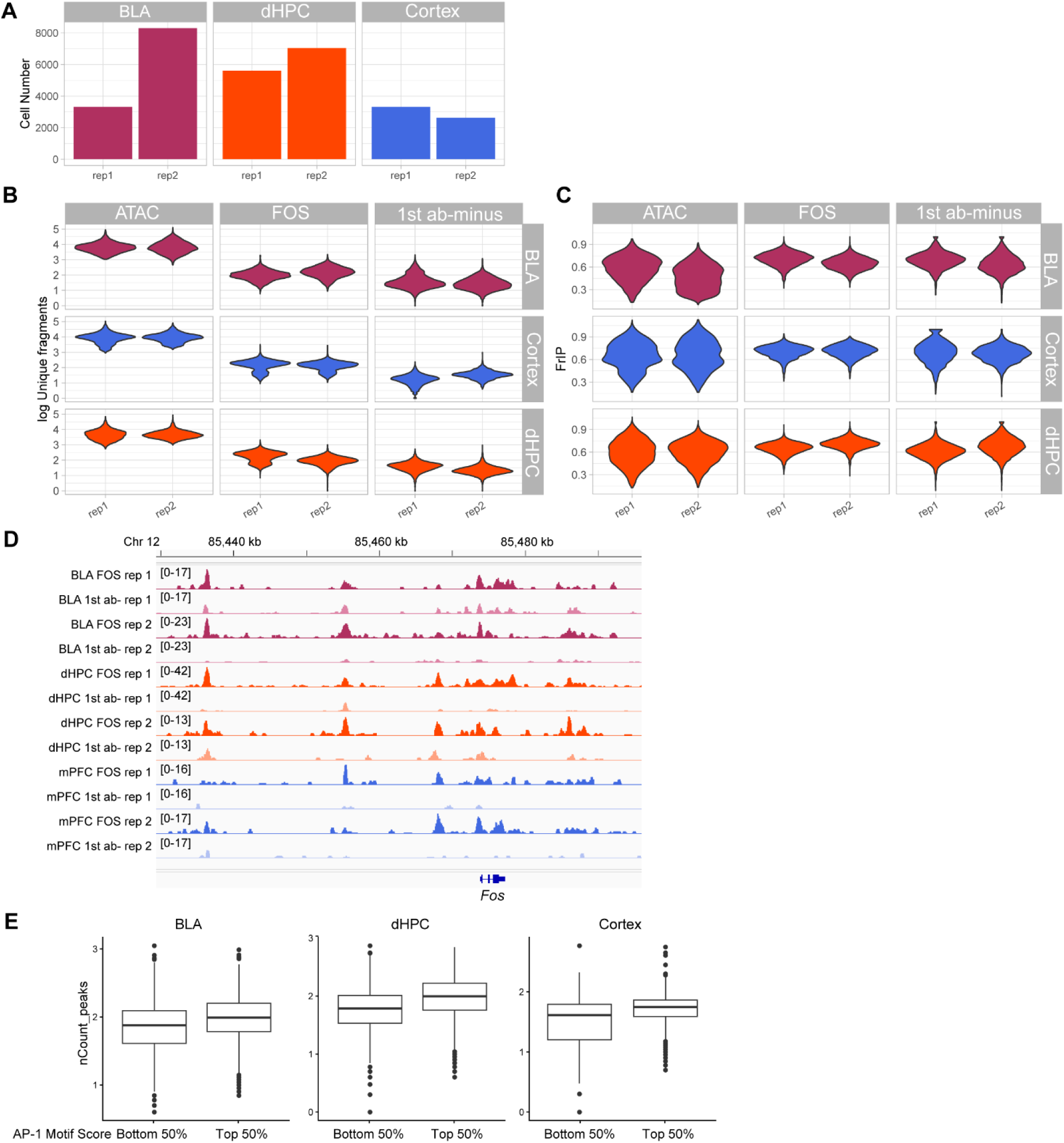
Quality control of FOS multi-nanoCTF. (A-C) Quality control metrics for multi-nanoCTF, including recovered cell numbers (A), unique fragments (B) and FRiP scores (C). (D) Genome browser views of multi-nanoCTF signals near *Fos* locus. FOS occupancy and first antibody-minus background are shown. (E) Box plots showing FOS-modality fragment counts within ATAC-defined peaks in nuclei stratified into top and bottom 50% groups by ATAC-based AP-1 motif score within each brain region. AP-1 motif scores were positively associated with FOS multi-nanoCTF signals.

**Supplementary Figure 6.**
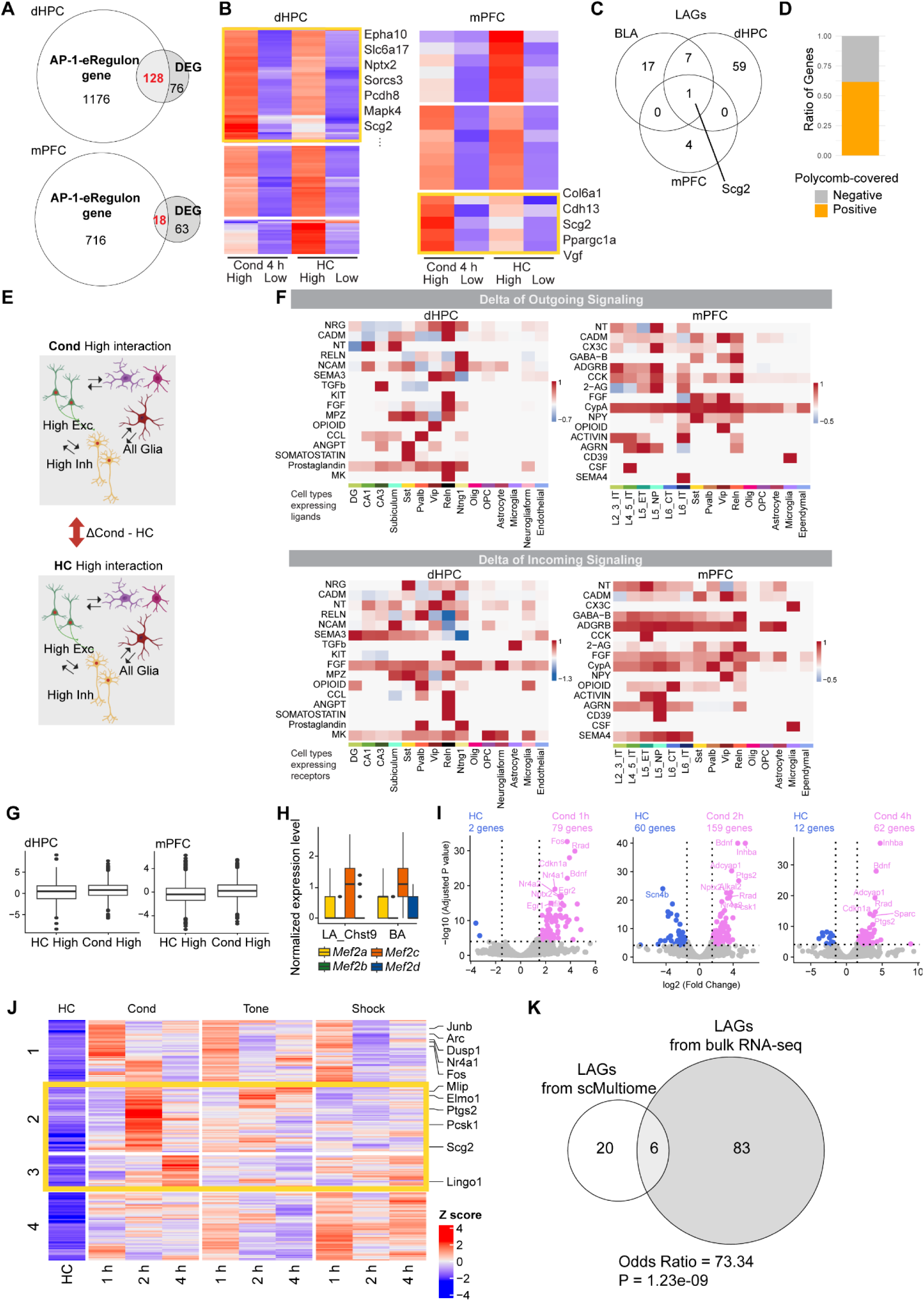
Characterization of learning-associated genes (LAGs). (A) Venn diagrams showing overlaps between ChromTRAP-associated DEGs and AP-1 eRegulon genes in dorsal hippocampal and mPFC excitatory neurons. Overlapping genes were defined as AP-1-associated secondary response genes (AP-1-SRGs). (B) Heatmaps showing hierarchical clustering of AP-1-SRG expression in dorsal hippocampal and mPFC excitatory neurons. Yellow boxes indicate learning-associated genes (LAGs). (C) Venn diagram showing overlap of LAGs across brain regions. (D) Bar plot showing the fraction of Polycomb-covered LAGs in BLA. (E) Schematic of CellChat comparison between Cond High and HC High populations, while retaining glial populations as potential interaction partners. (F) Heatmaps showing differences in outgoing and incoming CellChat-inferred signalling strength between Cond High and HC High populations. All molecular interactions detected in either Cond High or HC High populations were visualized, revealing overall stronger signalling in Cond High and suggesting learning-associated enhancement of intercellular molecular interactions in putative engram cells. (G) Box plots showing MEF2 motif scores in HC High and Cond High neurons in dorsal hippocampal and mPFC excitatory neurons. (H) Box plot showing normalized expression of MEF2 family genes in BLA excitatory neurons. (I) Volcano plots showing differentially expressed genes in BLA FOS+NeuN+ nuclei 1, 2 and 4 h after conditioning compared with HC NeuN+ nuclei. Upregulated genes are shown in pink, downregulated genes in blue and non-significant genes in grey. (J) Heatmap showing expression patterns of genes upregulated after conditioning across HC, tone-shock conditioning, tone-only and shock-only groups. Genes were clustered based on their temporal expression patterns after conditioning. Yellow boxes indicate clusters of 89 genes preferentially induced by tone-shock conditioning. (K) Venn diagram showing overlap between LAGs identified by ChromTRAP-based analysis of scMultiome and conditioning-associated genes identified by bulk RNA-seq. Statistical significance was assessed by one-sided Fisher’s exact test.

